# Genomic Landscape of Patients with Germline *RUNX1* Variants and Familial Platelet Disorder with Myeloid Malignancy

**DOI:** 10.1101/2023.01.17.524290

**Authors:** Kai Yu, Natalie Deuitch, Matthew Merguerian, Lea Cunningham, Joie Davis, Erica Bresciani, Jamie Diemer, Elizabeth Andrews, Alice Young, Frank Donovan, Raman Sood, Kathleen Craft, Shawn Chong, Settara Chandrasekharappa, Jim Mullikin, Paul P. Liu

## Abstract

Germline *RUNX1* mutations lead to familial platelet disorder with associated myeloid malignancies (FPDMM), which is characterized by thrombocytopenia and a life-long risk (35-45%) of hematological malignancies. We recently launched a longitudinal natural history study for patients with FPDMM at the NIH Clinical Center. Among 29 families with research genomic data, 28 different germline *RUNX1* variants were detected. Besides missense mutations enriched in Runt homology domain and loss-of-function mutations distributed throughout the gene, splice-region mutations and large deletions were detected in 6 and 7 families, respectively. In 24 of 54 (44.4%) non-malignant patients, somatic mutations were detected in at least one of the clonal hematopoiesis of indeterminate potential (CHIP) genes or acute myeloid leukemia (AML) driver genes. *BCOR* was the most frequently mutated gene (in 9 patients), and multiple *BCOR* mutations were identified in 4 patients. Mutations in 7 other CHIP or AML driver genes (*DNMT3A, TET2, NRAS, SETBP1, SF3B1, KMT2C*, and *LRP1B*) were also found in more than one non-malignant patient. Moreover, three unrelated patients (one with myeloid malignancy) carried somatic mutations in *NFE2*, which regulates erythroid and megakaryocytic differentiation. Sequential sequencing data from 19 patients demonstrated dynamic changes of somatic mutations over time, and stable clones were more frequently found in elderly patients. In summary, there are diverse types of germline *RUNX1* mutations and high frequency of somatic mutations related to clonal hematopoiesis in patients with FPDMM. Monitoring dynamic changes of somatic mutations prospectively will benefit patients’ clinical management and reveal mechanisms for progression to myeloid malignancies.

**Key Points:**

- Comprehensive genomic profile of patients with FPDMM with germline *RUNX1* mutations.
- Rising clonal hematopoiesis related secondary mutations that may lead to myeloid malignancies.

## Introduction

RUNX1 is a transcription factor indispensable for the development and function of definitive hematopoietic stem cells (HSCs)^1^. Chromosome translocations and somatic mutations affecting *RUNX1* are frequently detected in hematologic malignancies, such as myelodysplastic syndrome (MDS), acute myeloid leukemia (AML), and acute lymphoblastic leukemia (ALL)^2^. Germline *RUNX1* mutations lead to familial platelet disorder with associated myeloid malignancy (FPDMM, OMIM #601399), a rare autosomal dominant disease associated with platelet defects resulting in easy bleeding and bruising^3^. Patients with FPDMM are predisposed to hematologic malignancies^4,5^, with a 35-45% lifetime risk of developing MDS, AML, chronic myelomonocytic leukemia, and ALL^6–8^. The current consensus for the incomplete penetrance of malignancy is that germline *RUNX1* mutations are insufficient for leukemogenesis. Additional risk factors, such as somatic mutations, are important for the development of hematologic malignancies^4,9^.

Somatic mutations in HSCs may lead to accelerated proliferation and reduced cell death, resulting in clonal expansion of the mutation-carrying HSC, or clonal hematopoiesis (CH) ^10,11^. CH increases with age. Large population studies discovered that CH increases the risks of atherosclerotic cardiovascular diseases^12,13^, hematologic neoplasms^11,14^, and other nonmalignant diseases^15^. Previous studies have described early-onset of CH in non-malignant patients with FPDMM^4,16^. However, previously studied cohorts were relatively small to find enough correlation between genomic alterations and malignant progression. Moreover, previously published genomic data were not associated with sufficient phenotypic data to confirm the role of CH in the development of hematologic malignancies.

To improve our understanding of FPDMM pathogenesis and to identify potential driver alterations for malignancy transformation, we initiated a natural history study in 2019 to longitudinally investigate the genomic and clinical profile of FPDMM. Here, we report the genomic data from 66 enrolled patients in our study whose samples have been sequenced for research purposes by the end of 2021.

## Methods

### Patients and samples

Patients were enrolled in the clinical study entitled “Longitudinal Studies of Patients with FPDMM” (ClinicalTrials.gov identifier: NCT03854318), after informed consent in accordance with the Declaration of Helsinki. *RUNX1* variants were determined to be pathogenic (P), likely pathogenic (LP), or variants of uncertain significance (VUS) by ACMG criteria^17^ and only those with P or LP variants were included in this study. Clinical studies of the enrolled participants are described in the accompanying clinical report by Cunningham et al.

Genomic DNA, RNA, and cryopreserved cell samples were processed and biobanked for further needs.

### Exome Sequencing and data processing

Exome sequencing (ES) of genomic DNA samples at the NIH Intramural Sequencing Center (NISC) is described in Supplemental Methods. Detailed information about the sequenced samples is described in Supplemental Table 1 and Supplemental Figure 1. Sequencing data were analyzed with in-house pipelines on NIH high-performance computing system “Biowulf”. Detailed descriptions of data analysis, workflow and parameters can be found in Supplemental Figure 2 and Supplemental Methods.

### Bulk RNA sequencing and analysis

RNA sequencing was performed at NISC with Illumina TruSeq strategy and PE151 chemistry on Novaseq 6000. The sequenced samples are listed in Supplemental Table 1 and detailed information on sequencing and data analysis is described in Supplemental Methods.

### Cytogenetics and CNV analyses

Cytogenetic analyses of bone marrow cells were conducted at the Mayo Clinic Laboratories. For single-nucleotide polymorphism (SNP) array, Infinium OmniExpressExome-8 kit was used to analyze gDNA samples (Supplemental Table 1). CNVPartition and PennCNV^18^ were used to identify candidate CNV events. CNVkit^19^ (v.0.9.8) was used to identify CNVs from ES data for samples without SNP array data. All CNV calls were revised with IGV illustration.

### Data availability

The raw ES, RNA-seq and SNP-array data have been deposited to dbGaP database under accession number phs003075.

## Results

### Study cohort and *RUNX1* variant evaluation

The natural history study was launched in early 2019 and by the end of 2021, 129 patients and 85 family controls had been enrolled. This report focuses on 66 patients in 29 families whose research genomic data are available. Sequential samples have been collected for 19 patients who have visited NIHCC more than once (Figure 1A, Supplemental Table 1).

**Figure 1.**
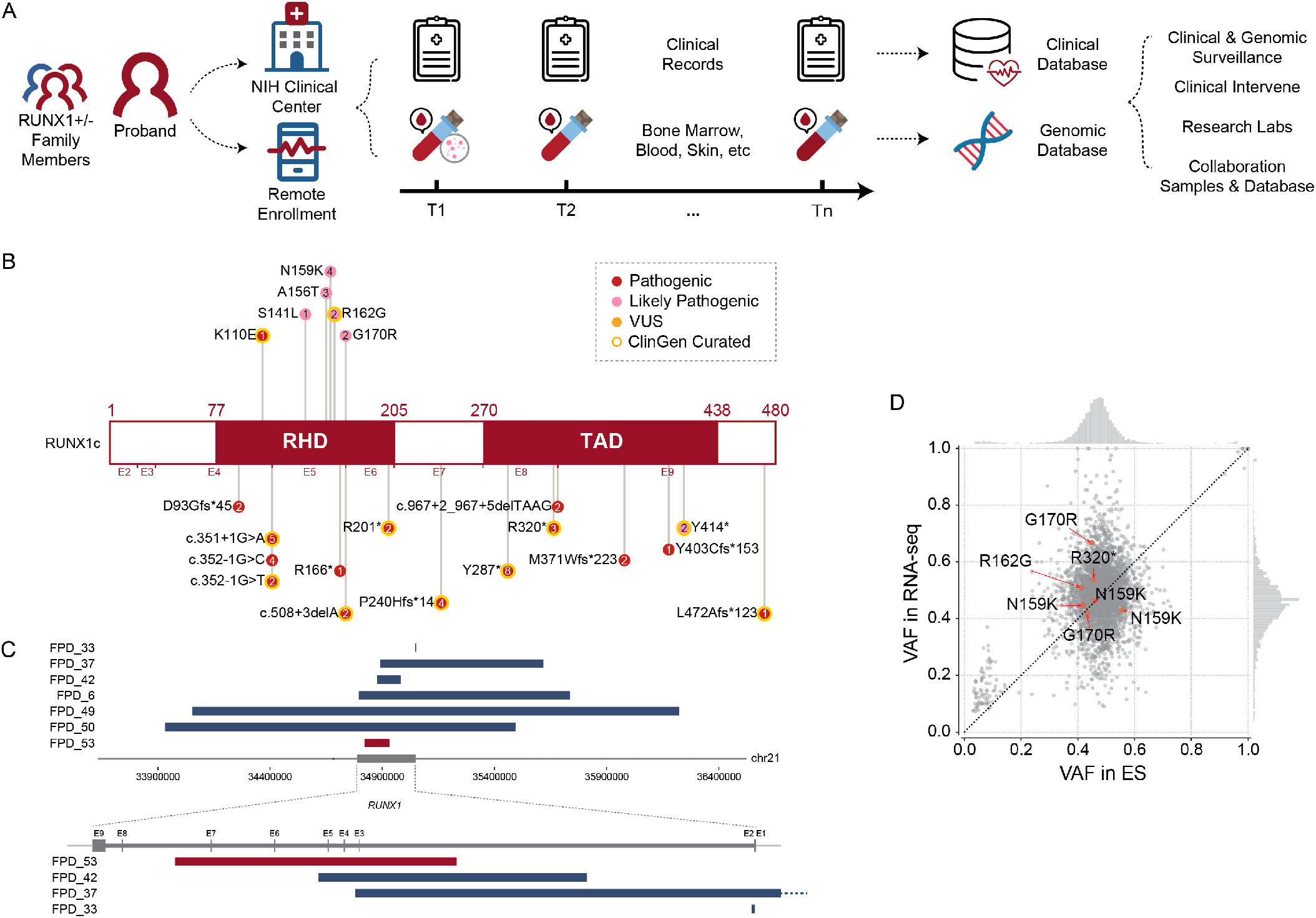
Overview of NIH FPDMM natural history study. (A) Study schema. Enrolled patients with germline *RUNX1* variants and family member controls visit NIH clinical center and/or send remote samples annually. Both clinical and research data stored in the clinical and genomic databases are used for downstream research and guide clinical management. (B) *RUNX1* mutations carried by the patients included in this manuscript. Numbers in the circle indicate patient number that carry the mutation. (C) CNVs affecting RUNX1 gene. Blue bars are showing deletions and red bars are showing duplication. (D) VAF correlation between exome sequencing and RNA-seq data. Red dots are *RUNX1* variants and grey dots are all other variants in all samples. Histograms on the top and right sides of the panel show the VAF distributions in exome sequencing and RNA-seq, respectively.

In total, 28 different germline *RUNX1* variants were detected in the 29 families. The most common types of *RUNX1* mutations are loss-of-function (LoF) mutations (23 families), including splice-site mutations, frameshifting indels, stop-gain mutations, and deletions that cause complete or partial loss of the *RUNX1* gene (Figure 1B and 1C, and Supplemental Table 2). Six families have 6 different missense mutations; all located in the Runt homology domain and predicted to be pathogenic or likely pathogenic per ACMG criteria^17^.

Large scale genomic alteration is common in our study cohort. Among the 29 families, three have large deletions covering the entire *RUNX1* gene, and in some cases additional genes. Three other families have smaller deletions, and one family has an intra-genic duplication that affects several *RUNX1* exons (Figure 1C).

To determine if the mutations alter *RUNX1* expression, we compared allele frequencies between ES and RNA-seq data from 7 patients, which have adequate coverage. The *RUNX1* variant alleles were expressed between 40%-70% at the RNA level, similar to VAF distributions of germline variants in other genes (Figure 1D).

### *RUNX1* splice site mutations

Multiple families have mutations at or near splice donor or acceptor sites, located in introns 4, 5, and 8 (Figure 1B). These mutations led to aberrantly spliced transcripts as detected by RNA-seq (Figure 2, Supplemental Figure 3). The c.351+1G>A variant, detected in two independent families, caused two types of exon 4 skipping: E2-E5 and E3-E5 (Figure 2B, Supplemental Figure 3B). Junction counts showed a significantly higher proportion of the novel splicing products than wildtype. The c.352-1G>C variant led to the usage of a cryptic acceptor site in exon 5 (Figure 2C, Supplemental Figure 3C), which is also the cryptic acceptor site resulting from a c.352-1 G>T mutation^3^. In this case, the mutant and wild-type transcripts were detected at similar levels. A cryptic splice donor site near the end of exon 5 was identified in c.508+3delA patients (Figure 2D, Supplemental Figure 3D), as previously reported^20^, and the aberrant splicing product count was less than that of the wildtype. Interestingly, a missense variant c.508G>C (p.G170R) affects the adjacent splice donor site at the end of exon 5, leading to the usage of the same cryptic splice donor site associated with c.508+3delA (Figure 2E, 2H, and Supplemental Figure 3D). For both variants, the splice product resulting from the cryptic donor site had fewer counts when compared to that of the wildtype junction (Supplemental Figure 3D). Finally, a c.967+2_967+5delTAAG variant caused exon 8 skipping (Figure 2G and I, type 7) and the activate of cryptic splice donors in intron 8 (Figure 2G and I, types 5 and 6), one of them (type 5) has been reported previously ^21^. At the protein level, it is predicted that the first five splice-site related variants will produce truncated RUNX1 proteins, while the c.967+2_967+5delTAAG variant will lead to two in-frame products: an insertion of 37 amino acids in the middle of TAD domain for type 6, and a deletion of 55 amino acids in the beginning of TAD domain for type 7 (Figure 2J).

**Figure 2.**
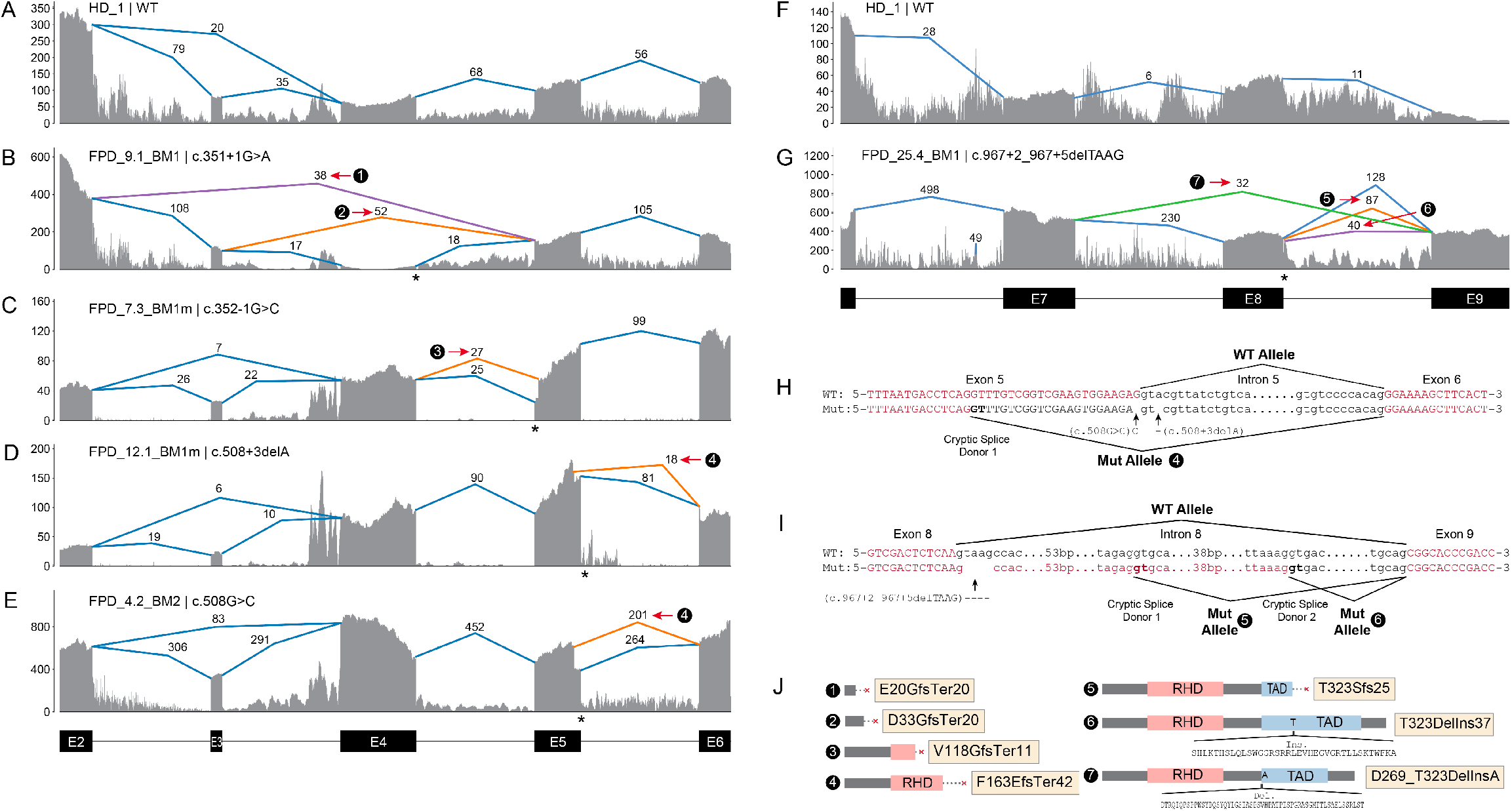
Aberrant splicing events in patients with *RUNX1* splice-site variants. (A) Splice junctions and covered reads from exons 2 to 6 of *RUNX1*, in a heathy donor bone marrow sample. (B-E) Splice junctions and covered reads in four patients with different *RUNX1* splice-site or splice region mutations. Orange or purple lines indicate abnormal junctions and blue lines indicate junctions also seen in the healthy control. Numbers indicate supporting read counts. (F) Splice junctions and covered reads from exons 6 to 9, in a heathy donor bone marrow sample. (G) Splice junctions and covered reads in a patient with c.967+2_967+5delTAAG mutation. (H, I) Detailed information of cryptic splice donors and abnormal splice junctions caused by c.508+3delA, c.508G>C and c.967+2_967+5delTAAG. (J) Predicted protein products of the six abnormal splicing transcripts shown in panels B-E and G.

### Somatic mutation landscape in patients with FPDMM

We have generated ES data from 62 patients with FPDMM for somatic mutation identification. We applied two strategies (Supplemental Figure 2) to identify somatic mutations. For 31 patients with ES data from fibroblast, true somatic mutations were confirmed by comparing BM/PB data with fibroblast data. For the remaining 31 patients without fibroblast data, we identified likely-somatic mutations in the PB or BM samples based on their absence in family members, as well as their population frequency and presence in the Catalogue Of Somatic Mutations In Cancer (COSMIC) database. The number of likely-somatic mutations in these patients is likely an underestimate since we have been conservative with somatic mutation calling.

The somatic mutation landscape of hematopoietic cells in the patients with FPDMM of our cohort is depicted in Figure 3A. The middle heatmap shows the aggregated somatic mutation landscape that merged all mutations in CHIP or AML driver genes (CL genes) detected in each patient. Interestingly, 24/54 (44.4%) of non-malignant and 6/8 (75%) of patients with hematologic malignancy have at least one somatic mutation in CL genes (Supplemental Table 3). Somatic mutations were recurrently (>1 patient) seen in the following CL genes: *BCOR, TET2, DNMT3A, KRAS, LRP1B, SF3B1, KMT2D, KMT2C, SETBP1, IDH1, NRAS*, and *PHF6. BCOR* was the most frequently mutated CL gene as *BCOR* mutations were found in 17.7% (11/62) of the patients and most *BCOR* mutations resulted in frameshifts. Moreover, four patients had more than one somatic *BCOR* mutation detected at the same time. Recurrent mutations were also seen in 7 non-CL genes (*NFE2, PTPN14, RRBP1, GSTT1, PRKDC, SPTBN2*, and *KDM3A*).

**Figure 3.**
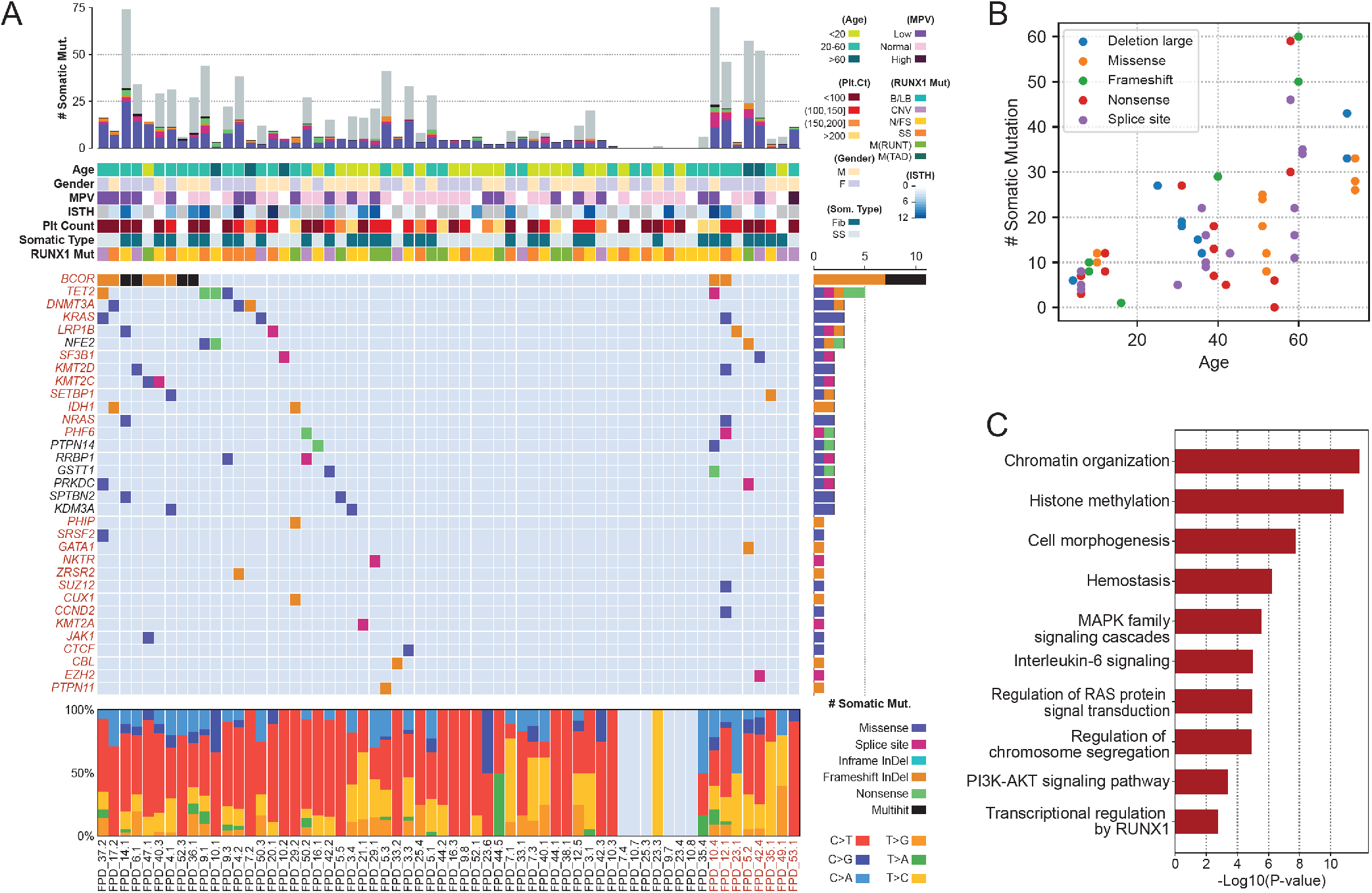
Somatic mutations of the NIH FPDMM cohort. (A) Somatic mutation landscape depicting CHIP genes, AML driver genes, and recurrently mutated genes. Each column is a patient; patients with hematologic malignancy are highlighted in red. The center heatmap shows somatic mutation distribution; mutation types are represented with specific colors, as described by legends at lower right. Each row is a gene; genes belonging to CHIP and AML driver genes are highlighted in red. The next heatmap above shows annotated demographic and clinical information, which are also color-coded as described by the legends on the right-hand side. The top bar plot shows the total somatic mutation numbers in each patient, with coding region mutations marked with colors used in the center heatmap and the non-coding region mutations colored in grey. The bar plot on the right of the center heatmap shows aggregated somatic mutation numbers for each gene, with color coding for mutation types. The bottom bar plot shows the percentage of the different base-changes (C>T, C>G, etc) in each patient. (B) Correlation between patients’ age at the time of sampling and the total number of somatic mutations. Only samples with fibroblast controls are included. The color code is the same as used in the center heatmap in panel A. (C) Functional enrichment analysis of all somatically mutated genes.

The overall somatic mutation numbers in each patient are shown in the top bar graph in Figure 3A. There seems to be a good correlation between the overall number of somatic mutations and the presence of CL gene mutations. Notably, 55.6% (30/54) of non-malignant patients had no mutation detected in CL genes. However, somatic CL mutations in patients without fibroblast ES data may have been under-detected (CL mutations were found in 19/31 patients with fibroblast ES data, while only in 11/31 patients without such data). As expected, the total numbers of somatic mutations correlated with patients’ ages (Figure 3B, Supplemental Figure 4B and 8B). Based on the published data from large cancer genome projects, myeloid malignancies show a relatively lower mutation burden than other cancer types^22,23^. For the 43 true-somatic non-malignant samples in our cohort, the median mutation burden is under 0.1 mutations/Mb, which is below the reported AML and MDS cohorts. Meanwhile, 10 samples with myeloid malignancies in our cohort showed mutation burden close to the TCGA-LAML cohort (Supplemental Figure 5A).

Between the top bar graph and the middle heatmap in Figure 3A are heatmaps for patient age, sex, mean platelet volume (MPV), the International Society on Thrombosis and Haemostasis Bleeding Assessment Tool (ISTH-BAT) score^24^, platelet count, as well as somatic mutation data type (whether fibroblast ES data is available) and *RUNX1* mutation type. The existence of any correlations between these measures and the detected somatic mutations is further depicted in Supplemental Figure 4, by sorting the landscape heatmap with different annotation items. Total somatic mutation number, CHIP mutations in genes most-frequently seen in the general population (*TET2, DNMT3A*), and in high-risk genes (such as *KRAS, NRAS, PHF6, ZRSR2*, and *SF3B1)* all trend up with increasing age (Supplemental Figure 4B). *BCOR* mutations were significantly enriched in patients between 20-60 years of age (Supplemental Figure 4B) and correlated with lower platelet count (Supplemental Figure 4D) and lower MPV level (Supplemental Figure 4E). Patients with multiple-hits in *BCOR* tend to have low platelet count (Supplemental Figure 4D). With current data, we did not find correlations between somatically mutated genes and *RUNX1* mutation types or gender.

The bottom bar plot of Figure 3A shows the types of base substitutions associated with the somatic mutations. C>T and T>C transitions are more common than transversions.

Pathway and gene ontology analyses of somatically mutated genes showed enrichment of regulation of histone methylation (also seen in CHIP studies^10,15^). Highly related pathways also include RAS, PI3K-AKT, MAPK, and IL6 signaling, which are related to inflammation (Figure 3C). In addition, mutations were enriched in genes with hemostasis functions and genes transcriptionally regulated by RUNX1.

### Recurrent somatic mutations in *NFE2*

As mentioned above, we found 7 recurrently mutated genes in the somatic mutation landscape besides the CL genes. Notably, *NFE2* was mutated in 3 unrelated patients, including two nonsense mutations and one missense mutation in the important BRLZ domain (Figure 4A). *NFE2* encodes a transcription factor involved in megakaryocyte development and platelet production^25,26^, which is mutated in more than 1% of MDS and around 0.7% of leukemia patients^27,28^ (Figure 4B). However, *NFE2* has not been reported as a gene associated with clonal hematopoiesis. Reported functional study also indicates that *NFE2* is a downstream target of RUNX1^29^. Published ChIP-seq data^30–32^ showed a strong RUNX1 binding signal in the promoter region of *NFE2* in CD34+ HSPCs (Figure 4C). In our cohort, 2 of the *NFE2* mutation carriers have already developed pre-monoclonal gammopathy of undetermined significance (pre-MGUS) and myeloma, respectively. Our data suggest that somatic *NFE2* mutations could contribute to FPDMM disease progression.

**Figure 4.**
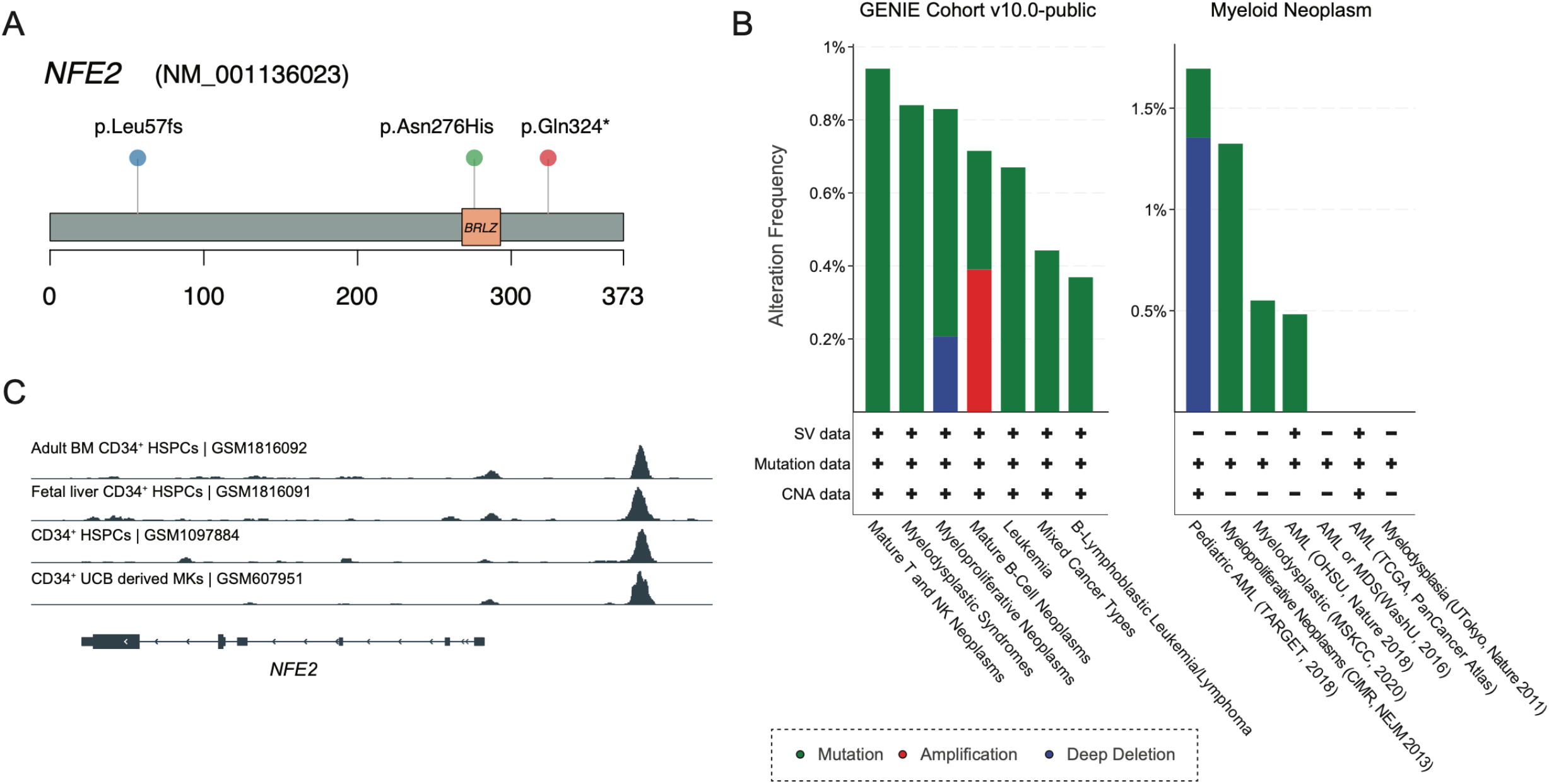
*NFE2* somatic mutations in FPDMM and hematologic malignancies. (A) Lollipop plot of *NFE2* somatic mutations in NIH FPDMM cohort. (B) Somatic alteration frequency of *NFE2* gene in AACR GENIE dataset and cBioportal myeloid neoplasm dataset. (C) CHIP-seq data on RUNX1 binding in NFE2 locus in CD34+ cells from four publicly available datasets^30–32^. Binding profile data were downloaded from CODEX database.

### Increased frequency of clonal hematopoiesis in patients with FPDMM

It has been reported that patients with FPDMM could develop detectable clonal hematopoiesis with a cumulative risk of >80% by 50 years of age^16^, which is far younger than the population average^33^, and that detection of these clones may help inform risk of developing hematologic malignancy^34^. We set out to determine the frequency of clonal hematopoiesis in our cohort by comparing our data with the population cohort reported by the TOPMed research program^35^. To meet the criteria applied to the TOPMed study, only somatic mutations detected in the previously reported 74 CHIP genes^35^ with VAF>5% were included for the comparison (Figure 5A, Supplemental Figure 6). 14/54 (26%) of the non-malignant patients in the FPDMM cohort have mutations in 11 CHIP genes at VAF>5%. This frequency is significantly higher than the general population (4.3%)^35^. Moreover, 13 of the 14 (92.9%) non-malignant patients with CHIP mutations are under 65 years of age; with median age of 42, and the youngest patient is only 13. In the general population, only 10% of people above 65 and 1% under 50 were reported to carry CHIP mutations. In our cohort, CHIP mutations were detected in 33.3% of patients (1/3) above 65, 18.8% of patients (9/48) under 50, and 4.3% of those (1/23) under 20.

**Figure 5.**
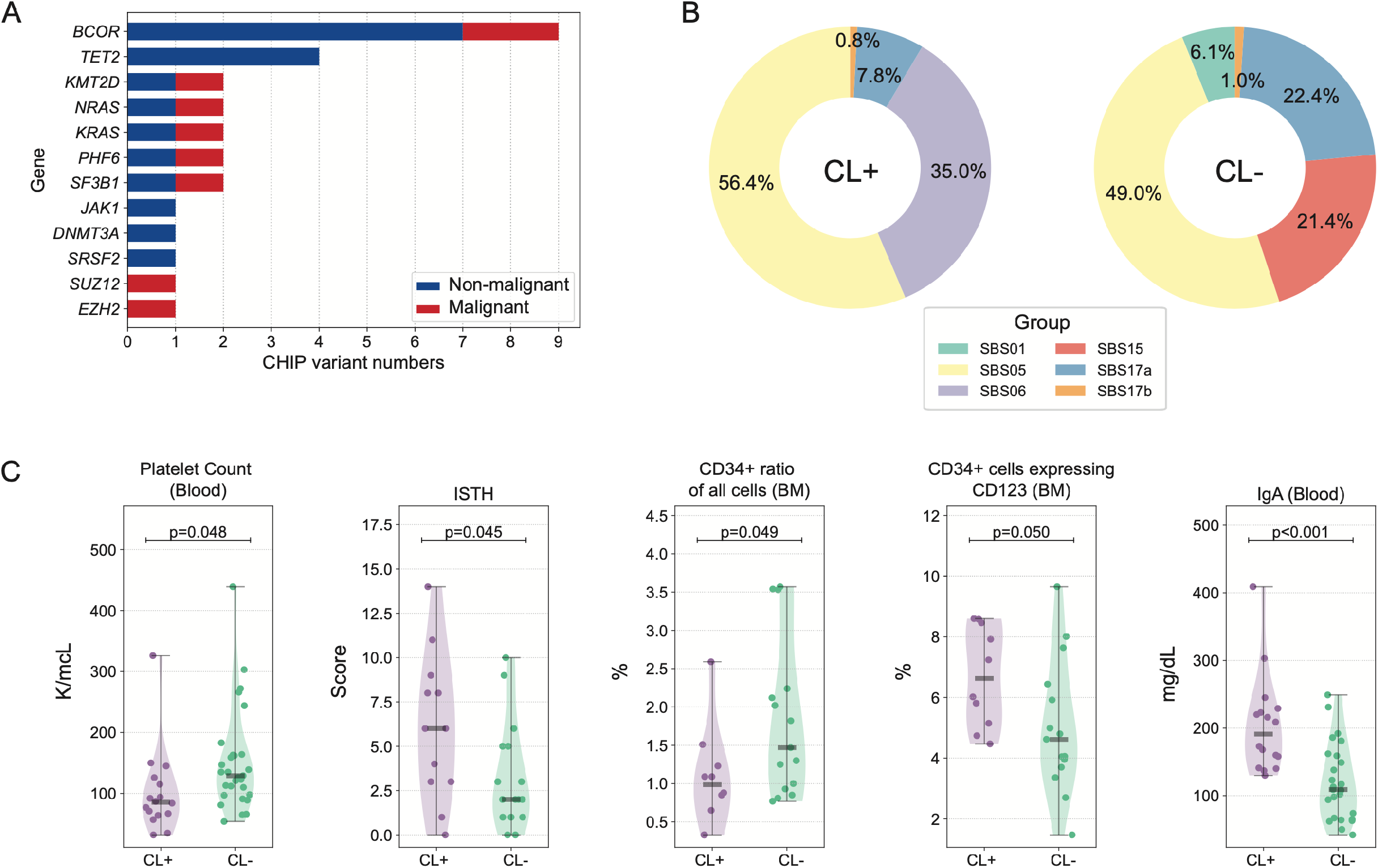
CHIP mutations in the NIH FPDMM cohort. (A) Bar graph showing the numbers of somatic mutations detected in 74 CHIP genes listed in the TOPMed study^35^. Only those with VAF>5% are included. (B) Mutation signatures^36^ in patients with or without somatic mutations in the 74-CL gene list. (C) Clinical phenotypes with significant difference between patient groups with or without somatic mutations in the 74-CL gene list. CL+: patients with somatic mutations in the CL genes. CL-: patients without somatic mutations in the CL genes.

We determined if there are differences in mutation signatures^36^ between patients with CL gene mutations and those without (Figure 5B, Supplemental Figure 5B and 5C). In both

CL+ and CL-groups, at least half of the mutations belonged to single-base substitution (SBS) signatures SBS1 and SBS5, which are both “clock-like” mutations^37^ that accumulate with time. Interestingly, in the CL+ group, 35% of the mutations were assigned to SBS6, which is associated with defective DNA mismatch repair^37,38^; in CL-group, 21.4% of the mutations are related to this, but classified as SBS15.

We also compared phenotype data between CL+ and CL-groups (Figure 5C). Patients in the CL+ group had significantly lower platelet counts (p=0.048) and higher ISTH-BAT scores (p=0.045) than patients in the CL-group. CL+ patients had lower numbers of CD34^+^ cells in the bone marrow (p=0.049), but a higher proportion of these cells also expressed CD123, which is overexpressed in many hematologic malignancies, including 80% of AML and B-ALL^39^. Moreover, CL+ patients had significantly higher blood IgA level (p<0.001), which may lead to increased pro-inflammatory cytokine production through activation of FcαRI-expressing immune cells^40,41^.

### Dynamic changes of somatic mutations over time

Multiple patients in our cohort have completed their 2^nd^ or 3^rd^ annual visits, and we have sequenced their samples to monitor the dynamic changes in their somatic mutations. We have observed a patient (patient #1) with a stable dominant clone characterized by a high VAF *BCOR* mutation (Figure 6A). Additional mutations in *DNMT3A* and *RUNX1T1* also remained stable at low VAF. Patient #2, who already developed MDS with ring sideroblasts, had a splicing factor *SF3B1* mutation which was stable at 25% VAF (Figure 6B). Mutations in *SF3B1* were reported in multiple chronic lymphocytic leukemia (CLL) and MDS patients^42^. However, VAFs of *THOC2* and *SMC2* mutations increased significantly at the second visit for patient #2 (Figure 6B). THOC2 is a member of the TREX complex, which is indispensable for mRNA export^43^; SMC2 is vital for the structural maintenance of chromosomes^44^. The combined annotation-dependent depletion (CADD) Phred-score of *THOC2* and *SMC2* mutations are 32 and 29.2, respectively, both predicting a highly deleterious risk. Similarly, we observed rising clones with potential risk in patient #3 (Figure 6C). In the 3rd yearly BM sample, we detected a new frameshift mutation in *ZRSR2*, which may cause dysregulated RNA splicing^45^, and a new somatic mutation in *IL6ST* which encodes a signal transducer shared by multiple cytokines, including IL6 and leukemia inhibitory factor^46^. Sequential data on more patients are shown in Supplemental Figure 7.

**Figure 6.**
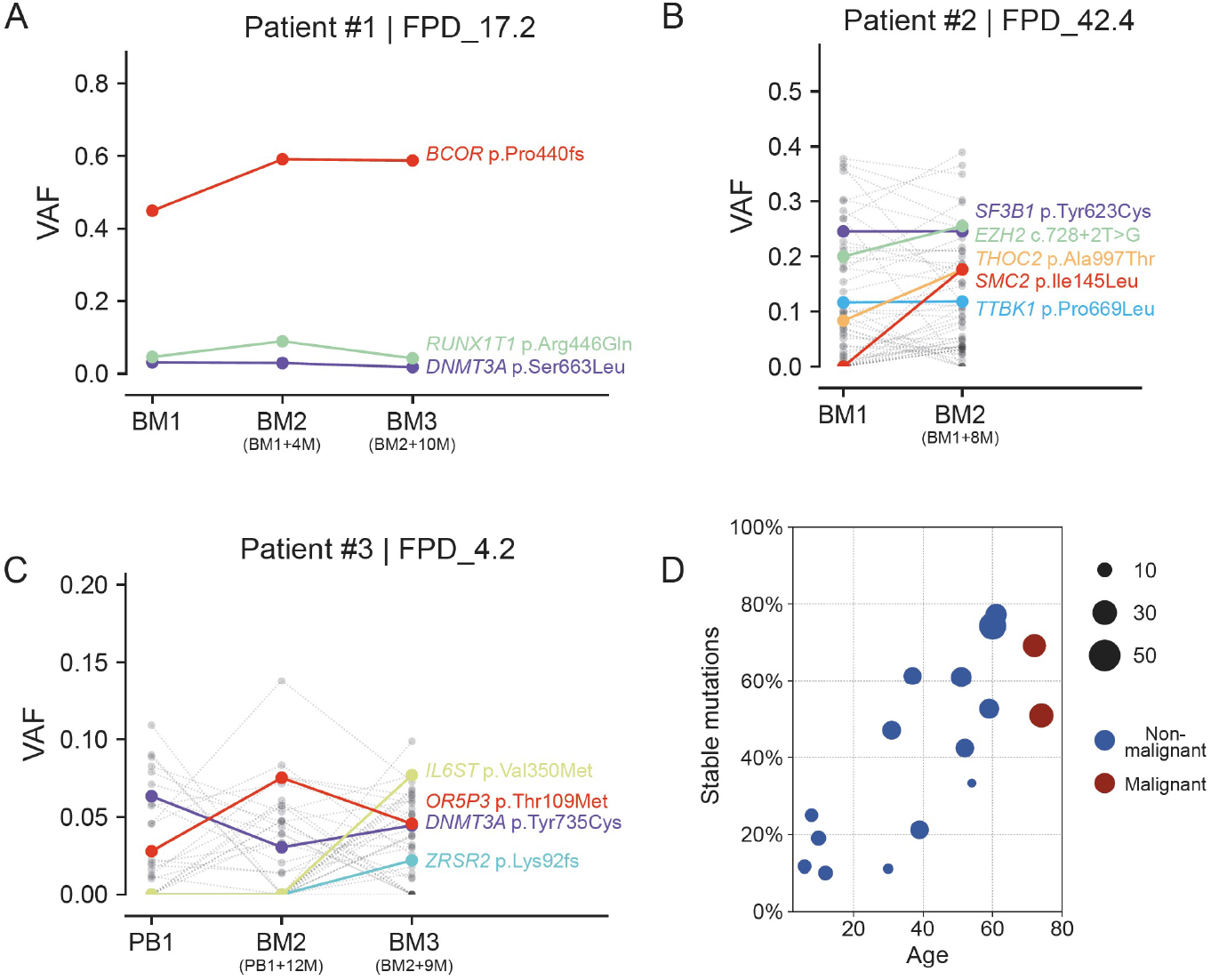
Somatic mutation VAF changes over time and age. (A-C) Somatic mutation VAF changes across samples collected at different timepoints. Colored dots and lines show mutations in CL and leukemia genes, grey dots and lines represent other somatic mutations. (D) Correlation between patient’s age and percentage of stable somatic mutations (present in at least 2 timepoints) in all somatic mutations. Dot size indicate the total somatic mutation number in this patient.

While comparing multi-timepoint mutations in our cohort, we observed a pattern that younger patients usually have fewer stable clones over time. As shown in Figure 6D and Figure S8A, fewer mutations were stable (present in at least 2 timepoints) in patients <30 years of age. On the other hand, the fraction of stable mutations increased with age, suggesting the presence of more stable clones in older patients.

### Additional genomic risk factors in FPDMM

Besides somatic mutations identified by ES, we also investigated other genomic alterations as potential risk factors cooperating with *RUNX1* mutations for malignant transformation.

With SNP-array-based copy number variation (CNV) analysis, we detected increased frequency of CNVs in patients with FPDMM. Specifically, we identified CNVs in 40% (10/25) of the non-malignant patients (Supplemental Table 4), while we detected CNV in only one of 8 family controls (12.5%). For most patients with CNV, there were no more than 2 CNVs, and only limited genomic regions were affected. By cytogenetics analysis, only one of the 51 analyzed non-malignant patients had a karyotype abnormality - a marker chromosome in the bone marrow cells (Supplemental Table 4). In addition, fusion-gene analysis did not reveal any in-frame fusion events in 21 patients for whom we have RNA-seq data.

We have also cataloged and analyzed germline variants in enrolled families. We focused on genes related to myeloid cell differentiation and hemostasis functions. Figure 7 demonstrates the distribution of the predicted deleterious germline variants in such genes in the enrolled families. We also included the percentage of patients in each family who have developed HMs, to highlight genes enriched in these high penetrance families. For example, germline *P2RX7* variants were detected in two families, one with a splice region variant and the other with a missense mutation, both are located in the P2X purinoceptor extracellular domain. In these two families, 71%, and 43% of *RUNX1* patients developed HMs, respectively. Previous studies reported that the *P2RX7*-related pathway could induce activation of tissue factor, which in turn may initiate thrombosis^47^. We also found *TNFSF9* variants in three families; the protein encoded by this gene belongs to the tumor necrosis factor (TNF) ligand family, and it correlated with platelet phenotypes in GWAS studies^48^. We found germline mutations in genes closely related to myeloid malignancies, such as *GATA2, PTPN11*, and *NF1*, and genes involved in biological functions related to the phenotype of FPDMM, such as *MMRN1, SERPINA10, SRF*, and *VWF*.

**Figure 7.**
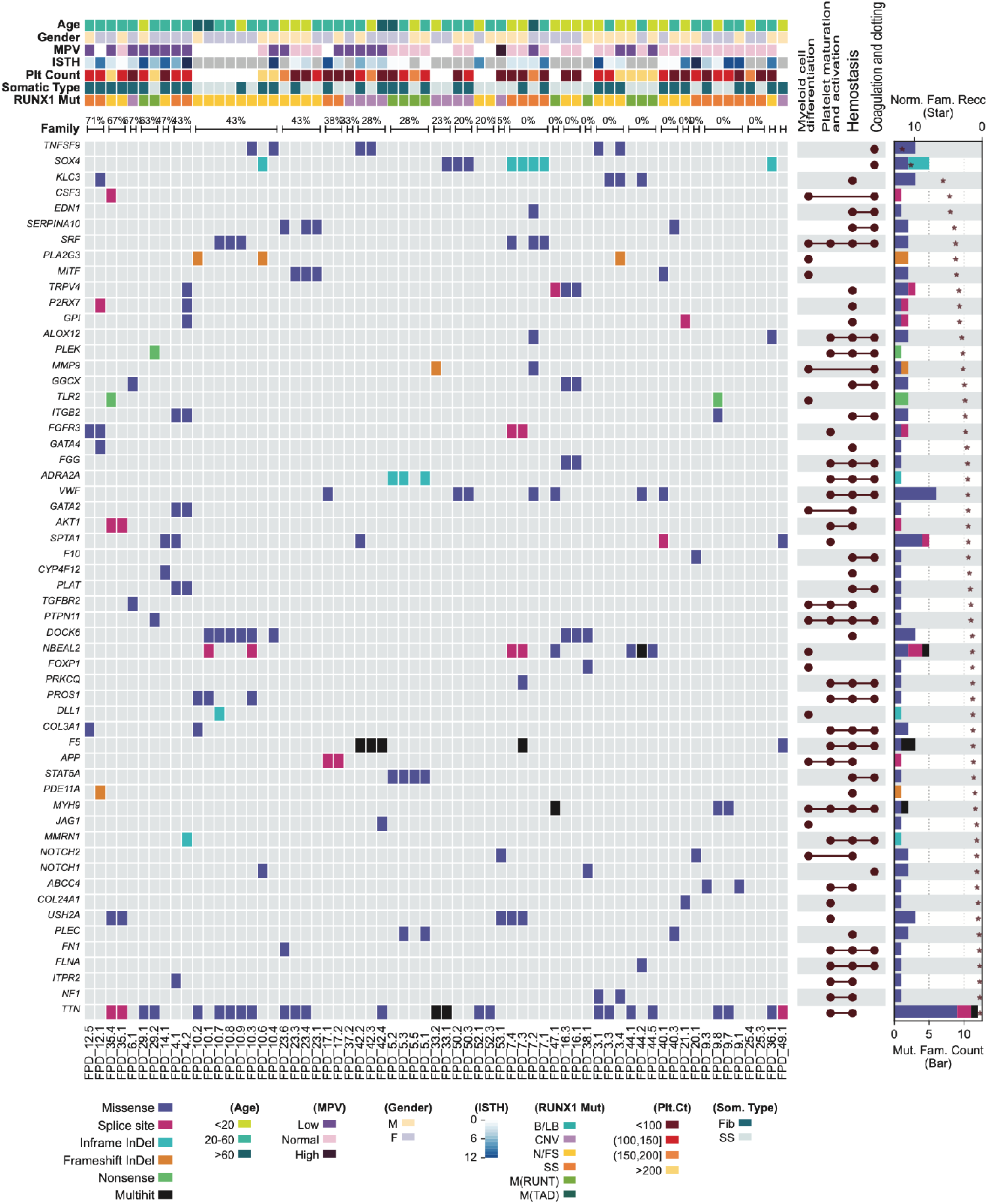
Heatmap of deleterious germline variants in genes related to myeloid cell development and hemostasis. Demographic and clinical data are depicted at the top, including percentages of affected family members who developed HM in each family. The UpSet plots on the right side of the heatmap show the related functional annotations for each gene. The bar plots on the far-right side show numbers of families that carry germline mutations in each gene. Protein-size-normalized family number are shown with star sign. The color codes for mutation types (both the heatmap and the bar plot) and for the clinical and demographic data are listed below the heatmap.

In our cohort, we identified predicted-deleterious germline variants in Fanconi anemia genes. Among the seven families with these variants, 4 of them showed a high rate of HMs within their family (Supplemental Figure 9A). Genes in the RTK-RAS-PI3K pathway also showed germline alterations (Supplemental Figure 9B). The exact significance of these germline variants is unknown; however, it is possible that stronger associations will be detected in the future through our longitudinal study.

## Discussion

In our cohort, germline variants were found to disrupt *RUNX1* mainly in two ways: (1) loss-of-function variants truncating or completely deleting the RUNX1 protein and (2) missense variants impacting critical functional domains. In addition, we have observed splice donor and acceptor sites as mutation hotspots in our patients, which lead to the usage of cryptic splice-sites and alternative splicing events, resulting insertions, deletions, frameshifts, and truncations. Due to the deleterious nature of the pathogenic germline *RUNX1* variants and the presence of mutational hotspot, therapeutic strategies can be envisioned to increase or restore *RUNX1* expression from the normal allele, and to block or repair hotspot mutations through antisense oligo, CRISPR or other targeted strategies.

Our somatic mutation landscape demonstrated that about half of the patients with FPDMM have at least one somatic mutation in a CL gene. Compared to patients without these mutations, the affected group showed significantly different phenotypes, including lower platelet count, lower percentage of CD34^+^ cells in the marrow, and higher blood IgA level. Functional enrichment analysis of the mutated genes identified several essential functions or pathways related to inflammation and RAS/PI3K-AKT-mTOR/MAPK. Mutation signatures also indicate that the affected group has a higher signal linked to DNA mismatch repair. Besides CHIP genes frequently seen in the general population, *BCOR* is the top mutated gene in patients with FPDMM^35^. Moreover, BM samples from four non-malignant patients harbored more than one *BCOR* mutation. In one of them, there were 5 different *BCOR* mutations, with VAFs ranging from 2.3% to 40%. From our data, most patients with *BCOR* mutations have not developed hematopoietic malignancy, even for cases who have *BCOR* mutations at relatively high VAFs. Therefore, the mutation mechanism and the impact of *BCOR* mutations on malignant transformation need further investigation. Overall, patients with FPDMM seem to have a distinct somatic mutation landscape, which is likely shaped by the pathogenic functions of mutated RUNX1, as recently described for Shwachman-Diamond Syndrome^49^.

Our study identified somatic mutations in several genes that were already reported in MDS or AML, such as *NRAS, KRAS, SF3B1, PHF6*, and *ZRSR2*. Interestingly, we found somatic *NFE2* mutations in 3 unrelated patients, including one who has developed myeloid malignancy. As a gene downstream of *RUNX1*, and involved in megakaryocyte development, *NEF2* has not received much attention in the context of FPDMM. It could be an important player and therefore a biomarker for developing malignancy.

We do not have enough evidence to connect other recurrently mutated genes with FPDMM. Histone demethylase KDM3A is an essential member of the JAK2-STAT3 signaling pathway^50^, which might be related to HM. AHI1 was reported as a member of AHI-1–BCR-ABL–DNM2 complex, which regulates leukemic properties of primitive CML cells^51^. Therefore, we will pay attention to these genes and further confirm their possible link to the disease development with additional data and through *in vitro* and *in vivo* functional studies.

In previous studies, somatic mutations in *RUNX1* were found to be common in patients with FPDMM who have developed malignancies^4^. However, for the 12 patients with HM in our cohort, only one carried a *RUNX1* somatic mutation - a large deletion. This frequency is far below the reported frequency (>40%). It is unclear why we have not seen more somatic mutations in *RUNX1* in our cohort. It would be interesting to see if we detect more *RUNX1* somatic mutations as we expand our cohort and continue our longitudinal study of the patients.

Longitudinal tracking somatic mutations can help us monitor dynamic changes of clonal expansion and transformation to malignancies. We have observed multiple patients with 2 or 3 timepoints, some of them have somatic mutations with stable VAFs, while others have mutations with fluctuating VAFs. Interestingly, younger patients tend to have higher VAF fluctuations between timepoints, while older patients tend to have more stable mutations. The clinical significance of these findings is presently unclear and hopefully will become more apparent with longer follow-up of these patients.

In addition to somatic mutations, deleterious germline variants in genes related to hematologic malignancies may also increase the risk of malignancy development in patients with FPDMM. However, determining which candidate germline variants are relevant to pathogenesis and progression of FPDMM is difficult without experimental validation or statistical power. We believe that after accumulating more data from the patients and their families, and performing experiments for functional confirmation, we will be able to identify germline modifiers that may stratify risk in patients with FPDMM.

Early and accurate detection of disease progression in FPDMM is imperative for clinical management and improving outcomes. We will continue following enrolled patients to collect more “snapshots” of their genomes and identify risk factors as early as possible. We hope to not only discover biomarkers for disease progression in FPDMM, but also develop improved clinical management strategies for all patients with FPDMM.

## Acknowledgements

We are grateful to patients and their family members for their participation in this clinical study. This work was supported by the Intramural Research Programs of the National Human Genome Research Institute and National Cancer Institute, NIH. The authors thank the NIH Intramural Sequencing Center (NISC), NHGRI Genomics, Microarray, Bioinformatics, Cytogenetics Core, and National Institute of Arthritis and Musculoskeletal and Skin Diseases (NIAMS) Genomic Technology Section for generating genomic data. We thank NIHCC for conducting clinical procedures, pathology evaluations, and lab tests. This work used the computational resources of the NIH HPC Biowulf cluster (http://hpc.nih.gov).

## Authorship Contributions

K.Y. and P.L. designed the analysis strategy and wrote the manuscript; K.Y., M.M., E.B., and J.Diemer performed the experiments; K.Y., R.S. analyzed the data; N.D. provided genetic counseling; J. Davis, N.D., M.M., and L.C. enrolled patients; M.M. and L.C. provided clinical care; L.C. served as medical director and P.L. served as protocol PI; all authors discussed and revised the manuscript.

