## Supplemental Methods for "Genomic Landscape of Patients with Germline *RUNX1* Variants and Familial Platelet Disorder with Myeloid Malignancy"

### Supplementary methods

#### Exome sequencing and data analysis

##### 1. Sequencing

Genomic DNA from patients' bone marrow, peripheral blood, and skin tissue-derived fibroblast were submitted to NIH Intramural Sequencing Center (NISC) for exome sequencing. In general, gDNA was fragmented with Covaris Focused-ultrasonicator into small fragments around 300 bp size. Swift Biosciences Accel-NGS 2S Plus DNA Library Kit was used to perform end repair, A-tailing, adapter ligation, and a limited cycle of amplification, following the standard protocol. 500 ng of each of the eight libraries were pooled together, following exome capture procedurals with IDT xGen Exome Research Panel. NimbleGen SeqCap EZ Hybridization and Wash Kit (Roche) were used for a few archived samples. Prepared libraries were sequenced on Illumina Novaseq 6000 platform with a paired-end strategy aiming to achieve 200X average depth. In addition, several archive data (EMKFPD samples) were captured with Roche NimbleGen SeqCap EZ Hybridization and Wash Kit and sequenced for about 80X average depth.

##### 2. Alignment and recalibration

Sequencing data were transferred to NIH high-performance computing system "Biowulf", and analyzed with an in-house pipeline. First of all, we used fastp[1] to do quality control of fastq files with default parameters. Low-quality reads and sequencing adaptor sequences were removed. Clean sequencing reads were mapped to hg38 reference with a streamed pipe of bwa[2] (v.0.7.17) alignment, sambamba[3] (v.0.8.0) sort bam files, and picard MarkDuplicates mark duplicates (v.2.17.11). Since most libraries were sequenced in multiple lanes, bam files of different lane's data were merged with sambamba. BQSR recalibration was further performed with GATK best practice[4] (v.4.2.0.0) to generate the analysis-ready bam file.

##### 3. Genotype analysis

First of all, we used GATK (v.4.2.0.0) to do joint genotyping. In brief, per sample genotype were identified by HaplogypeCaller module with parameters "--emit-ref-confidence GVCF --max-alternate-alleles 3 --read-filter OverclippedReadFilter", and validated by ValidateVariants module with parameters "-gvcf --validation-type-to-exclude ALLELES". All gvcf results of individual samples were used as input of GenomicsDBImport module, and followed by steps of joint genotyping. Including GenotypeGVCFs with parameters "-

G StandardAnnotation -G AS_StandardAnnotation --allow-old-rms-mapping-quality-annotation-data --merge-input-intervals"; and VariantFiltration module with parameters "--filter-expression "ExcessHet>54.69" --filter-name ExcessHet". We further performed variant recalibration for SNPs and InDels.

For InDels, recalibrate model were generated by VariantRecalibrator model with parameters "--trust-all-polymorphic -tranche 100.0 -tranche 99.95 -tranche 99.9 -tranche 99.5 -tranche 99.0 -tranche 97.0 -tranche 96.0 -tranche 95.0 -tranche 94.0 -tranche 93.5 -tranche 93.0 -tranche 92.0 -tranche 91.0 -tranche 90.0 -an FS -an ReadPosRankSum -an MQRankSum -an QD -an SOR -an DP --use-allele-specific-annotations -mode INDEL --max-gaussians 4 --resource:mills,known=false,training=true,truth=true,prior=12 --resource:axiomPoly,known=false,training=true,truth=false,prior=10 --resource:dbsnp,known=true,training=false,truth=false,prior=2", and the model was applied to InDels by ApplyVQSR module with parameters "--use-allele-specific-annotations --truth-sensitivity-filter-level 95.0 --create-output-variant-index true -mode INDEL".

For SNPs, we first generated model by VariantRecalibrator module with parameters "--trust-all-polymorphic -tranche 100.0 -tranche 99.95 -tranche 99.9 -tranche 99.8 -tranche 99.6 -tranche 99.5 -tranche 99.4 -tranche 99.3 -tranche 99.0 -tranche 98.0 -tranche 97.0 -tranche 90.0 -an QD -an MQRankSum -an ReadPosRankSum -an FS -an MQ -an SOR -an DP --use-allele-specific-annotations -mode SNP --output-model module.file.path --sample-every-Nth-variant 10 --max-gaussians 6 -resource:hapmap,known=false,training=true,truth=true,prior=15 -resource:omni,known=false,training=true,truth=true,prior=12 -resource:1000G,known=false,training=true,truth=false,prior=10 -resource:dbsnp,known=true,training=false,truth=false,prior=7". This model were feed as input for the second round VariantRecalibrator to generate VQSR model with parameters "--trust-all-polymorphic -tranche 100.0 -tranche 99.95 -tranche 99.9 -tranche 99.8 -tranche 99.6 -tranche 99.5 -tranche 99.4 -tranche 99.3 -tranche 99.0 -tranche 98.0 -tranche 97.0 -tranche 90.0 -an QD -an MQRankSum -an ReadPosRankSum -an FS -an MQ -an SOR -an DP --use-allele-specific-annotations -mode SNP --input-model module.file.path --max-gaussians 6 -resource:hapmap,known=false,training=true,truth=true,prior=15 -resource:omni,known=false,training=true,truth=true,prior=12 -resource:1000G,known=false,training=true,truth=false,prior=10 -resource:dbsnp,known=true,training=false,truth=false,prior=7". Finally, the VQSR model were used in ApplyVQSR module to generate the final VQSR calibrated SNPs. Variants with PASS tag and QUAL>100 were kept.

To identify possible missing variants, especially low VAF variants, we also used freebayes[5] (v.1.3.5) and LoFreq[6] (v.2.1.5) to do genotyping as supplementary results. In Freebayes analysis, default parameters were used, following filtration criteria including (QUAL>20, DP>8, QUAL/AO>10, SAF>0, SAR>0, RPR>1, RPL>1). LoFreq analysis was also used with default parameters, mutations didn't show in GATK, and freebayes results with PASS tag were kept.

Genotype matrix were combined from three software, then annotated with SNPEff[7] and SNPSift. Multiple filtration procedurals were used to filter unreliable or polymorphism. First, mutations were compared with a region filter. Mutations located in exclude region files were removed, including DAC list, DUKE list, LCR list, Homopolymer list, and Mappable issue list. In addition, only if variants located in the exome capture region with 150bp flank on both ends, 1000G mappable region, and GIAB mappable regions, the variants were kept. Second, we did VCF normalization for all the variants passed region filters, to ensure the parsimony and left alignment of variants. Third, the population frequency of 1000G project, ExAC database, and gnomAD3 database were used to annotate all variants. Variants with 1000G VAF<1% and ExAC VAF<1%, and gnomAD3 popmax<1% were kept. In the last, all variants were manually reviewed with IGVtools to minimize false positive results.

For all variants that passed all filters, we re-evaluated depth, VAF, DP4 counts, based on high-quality sequencing reads (sequence_length >= 80, not secondary_alignment, and mapping_quality>50), and generated the finalized genotype matrix.

#### 4. Somatic mutation calling

Somatic mutation identification is important to studying clonal hematopoiesis in FPDMM patients, and it's the key to discovering determinate factors of disease development. To improve the sensitivity of somatic mutation calling, we applied several software to identify somatic mutations independently and merged the mutation calling set together. To distinguish somatic mutations in bone marrow or peripheral blood cells, we collected skin biopsies from patients and expanded fibroblast, and further sequenced them as somatic control. However, since the fibroblast preparation procedure takes additional time, and it could fail, we have several patients who don't have skin fibroblast sequencing data ready for analysis. So, two somatic mutation analysis strategies were used in our work. For patients with fibroblast control data, we define the somatic mutations calling results as true somatic. For other patients without fibroblast control data, we have to use a single sample somatic mutation calling strategy. Since we cannot confirm the existence in other tissues, we define the somatic mutation calling set as likely somatic.

True somatic mutation calling strategy: Base Recalibrated bam files of bone marrow/peripheral blood sample and fibroblast sample were used as input for Mutect2[4] (v.4.2.0.0), strelka[8] (v.2.9.0), LoFreq[6] (v.2.1.5), and MuSE[9] (v.1.0).

For Mutect2, we used 10 family controls without *RUNX1* variant to generate panel of normal (PoN) for analysis. Mutect2 module took bam files, reference, panel of normal, gnomAD germline resource to detect somatic mutations, and generate orientation bias model file. The orientation bias model was further processed by LearnReadOrientationModel, GetPileupSummaries, and CalculateContamination with default parameters. Finally, the processed bias model was used to filter the somatic mutation list. Mutations that didn't pass the model were removed from the analysis.

To analyze somatic mutation with Strelka, paired bam files were firstly used by Manta (v.1.5.0) to identify candidate small indels. After that, paired bam files, and candidate small indels reported by Manta were used by Strelka to report somatic SNVs and InDels. Default parameters were used with both Manta and Strelka.

We also used LoFreq, and MuSE2 to identify somatic SNVs and InDels with default parameters. Finally, we queried the genotype matrix to compare the genotype of BM/PB sample to the fibroblast sample, and generated the merged true somatic mutation list.

Likely somatic mutation calling strategy: Mutect2 single sample mode was used to predict somatic mutation from BM/PB sample itself. Previous described panel of normal (PoN) was used in this step to compare with. Similar to sample-control Mutect2 analysis, LearnReadOrientationModel, GetPileupSummaries, and CalculateContamination were done and filtered the somatic mutations. Since we sequenced multiple affected and unaffected individuals from the same family, if the mutation can be found in any other family members, we treat it as germline variant.

#### 5. Somatic mutation filtering process

Similar to the genotyping process, mutations located in the region from DAC list, DUKE list, LCR list, Homopolymer list, and Mappable issue list were removed, we only kept mutations found in the exome capture region and flanking 150bp region in both upstream and downstream. In addition, to further remove mutations located in the low-complexity region, if the surrounding sequence met the criteria like: BBBBB, ABABABAB, ABCABCABC, ABCDABCD, mutations will be removed.

For the mutation evidence, the mutation locus must met "depth >=10, VAF>=0.03, and alt-allele-reads>=3" in somatic sample, and "depth >=8, VAF<0.01" in fibroblast control. To remove the strand-bias, the alt-allele-reads must include at least one supporting read from both chains. Because we would like to detect early events of somatic mutations in CHIP-related genes and leukemia driver genes, we called back all somatic mutations in these genes that failed the general filters, and reviewed the alignment carefully. If the somatic sample met "VAF>0.01 and alt-allele-reads>=3", and the alignment is clean, we kept these mutations in downstream analysis. However, to do the comparative analysis with the TOPMed study, we only included somatic mutations with VAF>0.05 in the provided 74 gene list from the TOPMed study.

##### 6. Other analysis.

We used Maftools to do the mutation burden analysis[10]. And used SigProfilerMatrixGenerator[11] to do the somatic mutation signature analysis. T-test was used to evaluate the significance in the genotype-phenotype analysis.

#### RNA sequencing and data analysis

##### 1. Sequencing

RNA was extracted from PB or BM samples collected with PAXgene Bone Marrow/Blood DNA Kit (Qiagen) or from isolated PB/BM mononuclear cells. We took 0.5-1 ug of total bone marrow or peripheral blood RNA as input of RNA-seq. The RNA quality must meet the RIN>7.

RNA-seq libraries were prepared by NIH Intramural Sequencing Center (NISC) with Illumina TruSeq Stranded Total RNA with Ribo-Zero Globin strategy, following by PE151 sequencing on Novaseq 6000 platform. In addition, libraries were sequenced on the Illumina Novaseq platform with PE151 chemistry.

##### 2. Analysis

Sequencing data were analyzed with an in-house pipeline under the snakemake pipeline framework. First of all, low-quality reads and sequencing adaptor sequences were removed by fastp with default parameters. STAR[12] (v.2.7.6a) was used to align all clean sequencing reads to human reference genome hg38, with two pass alignment strategy. Specific parameters were used in pass 1 (--outFilterMultimapScoreRange 1 --outFilterMultimapNmax 20 --outFilterMismatchNmax 10 --alignIntronMax 500000 --alignMatesGapMax 1000000 --sjdbScore 2 --alignSJDBoverhangMin 1 --outFilterMatchNminOverLread 0.33 --outFilterScoreMinOverLread 0.33), and pass 2 (--outFilterMultimapScoreRange 1 --outFilterMultimapNmax 20 --outFilterMismatchNmax

10 --alignIntronMax 500000 --alignMatesGapMax 1000000 --sjdbScore 2 --alignSJDBoverhangMin 1). Aligned bam files were then sorted and indexed with sambamba[3] (v.0.8.0). Splicing junction forms and counts were collected from the output of STAR.

We applied STARfusion[12] (v1.8.0) and Arriba[13] (v1.2.0) in the fusion gene analysis to maximize the detection of fusion events, followed the standard protocols.

#### SNP array tests and data analysis

For each sample, 300 ng high-quality genomic DNA was processed and loaded with OmniExpressExome-8 v1.6 bead chip. The data was processed with GenomeStudio, and CNV analysis was performed with two analysis plug-ins: cnvPartition and PennCNV[14]. We also illustrated the copy number Log R ratio (LRR) and B allele frequency (BAF) with an in-house script. All predicted CNV events were manually reviewed to eliminate false positive calls.

#### Other

We developed standardized exome genotyping, somatic mutation calling, and RNA-seq analysis pipelines with the snakemake framework[15]. All customized figures in this manuscript were generated with matplotlib.
