## Supplemental Figure for "Genomic Landscape of Patients with Germline *RUNX1* Variants and Familial Platelet Disorder with Myeloid Malignancy"

### Figure S1

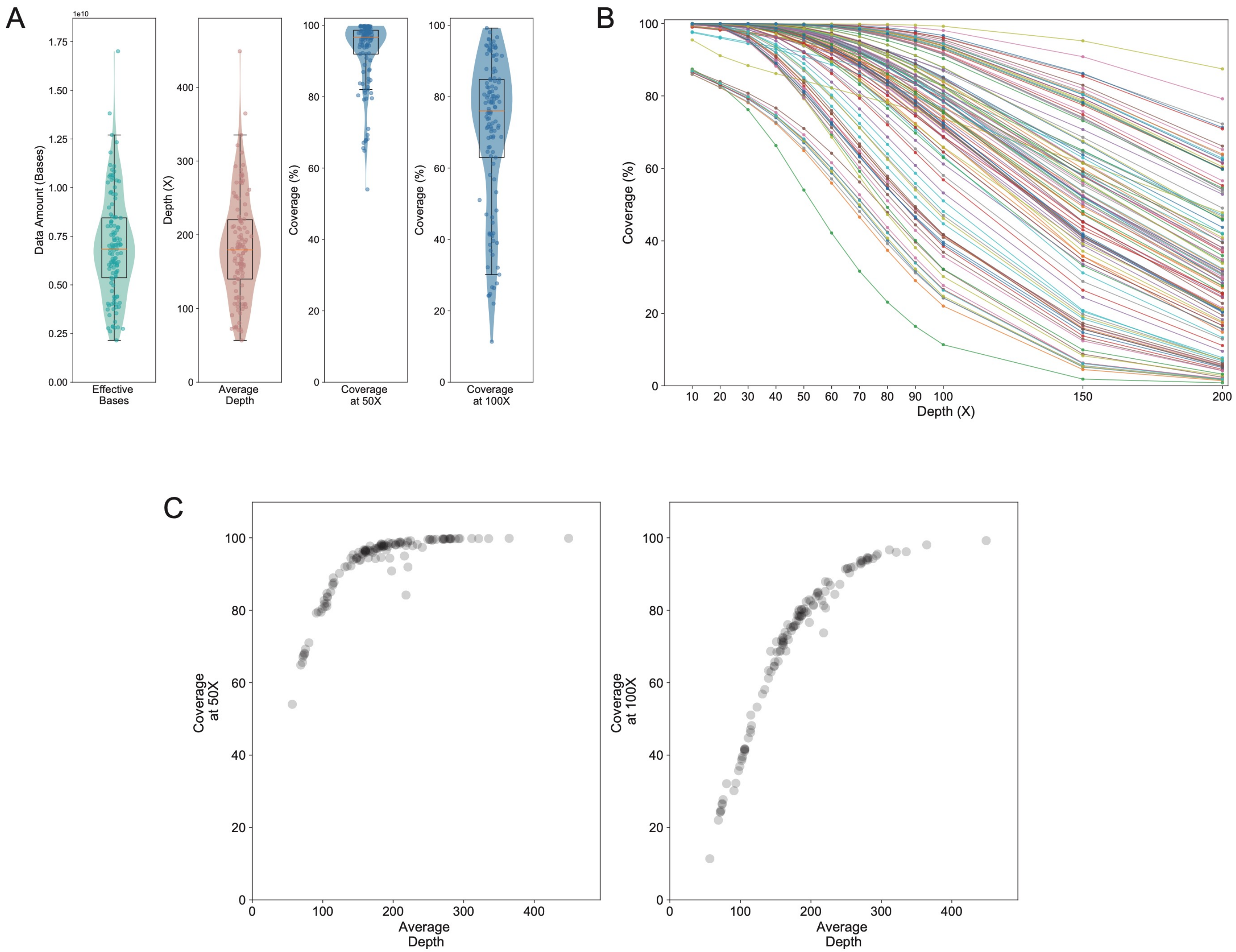

A
1e10
Data Amount (Bases)
Effective Bases
Depth (X)
Average Depth
Coverage (%)
Coverage at 50X
Coverage (%)
Coverage at 100X
B
Coverage (%)
Depth (X)
C
Coverage at 50X
Average Depth
Coverage at 100X
Average Depth

Figure S1. **Summary of exome sequencing data** (A) Distribution of effective sequenced bases, quantile-adjusted average depth, coverage of targeted region that covered at lease 50X, and coverage of targeted region that covered at least 100X. Each dot indicates a sequenced sample. (B) Distribution of exome coverage at specified minimum depth. (C) Sequencing saturation at 50X and 100X minimum depth.

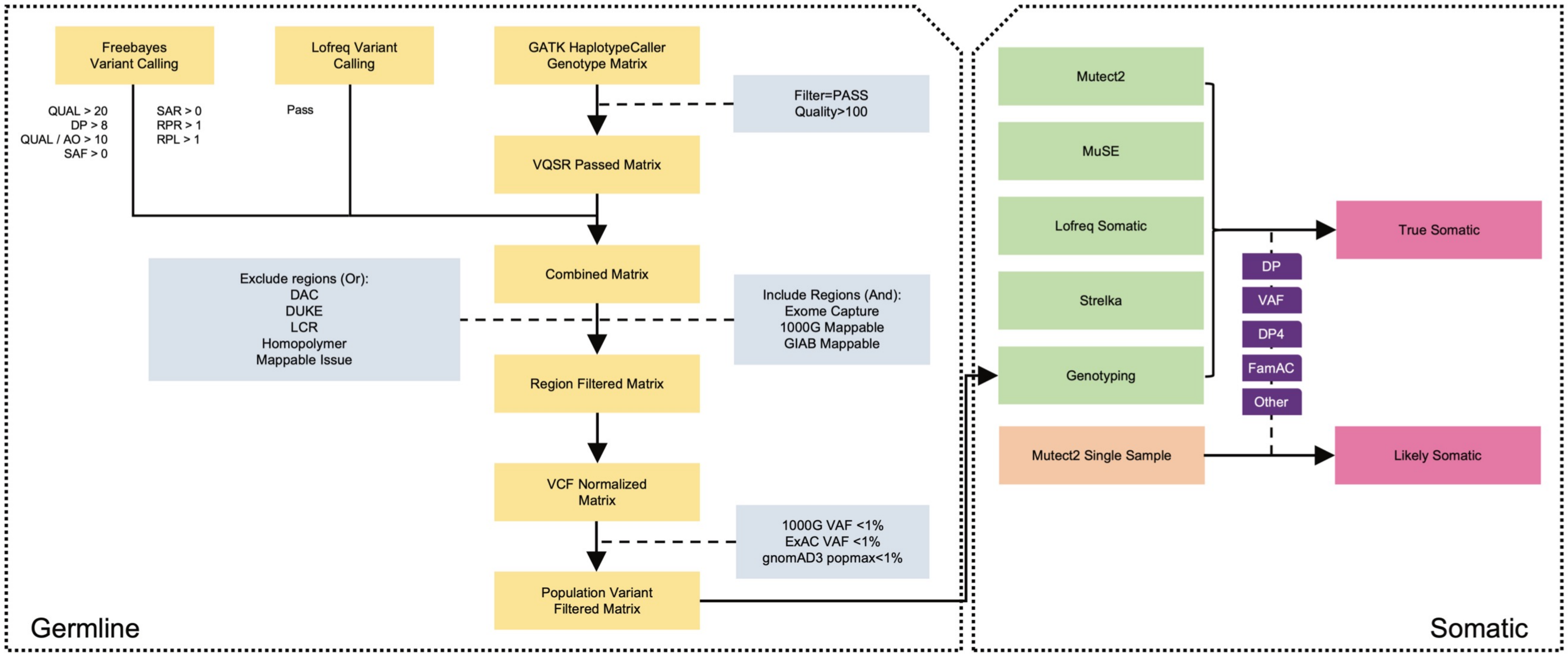


Figure S2. **Workflow of germline genotyping and somatic mutation calling**

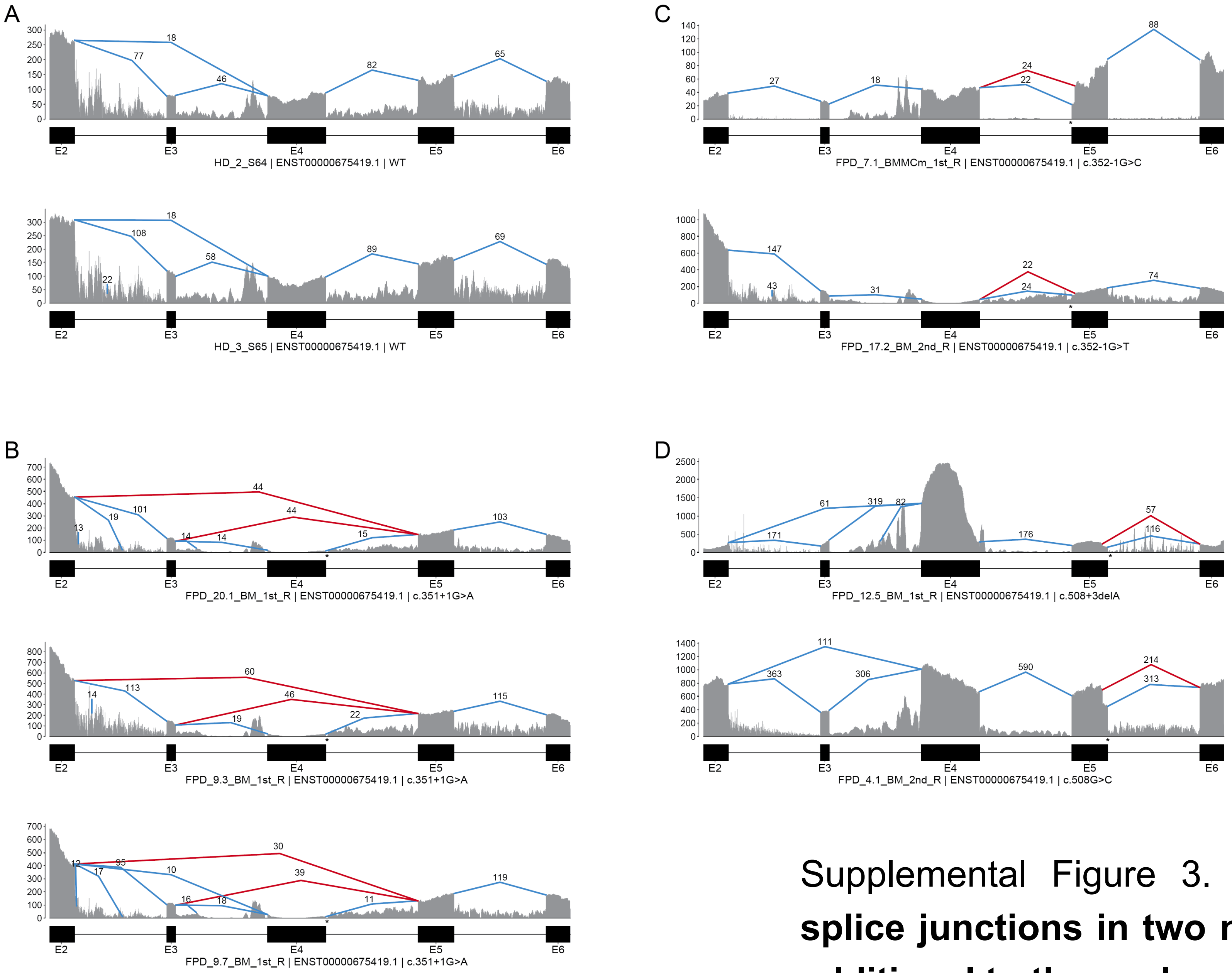


Supplemental Figure 3. **Junction plots showing *RUNX1* splice junctions in two normal controls and 7 patients (in additional to those shown in Fig. 2A-G)**

### Figure S4

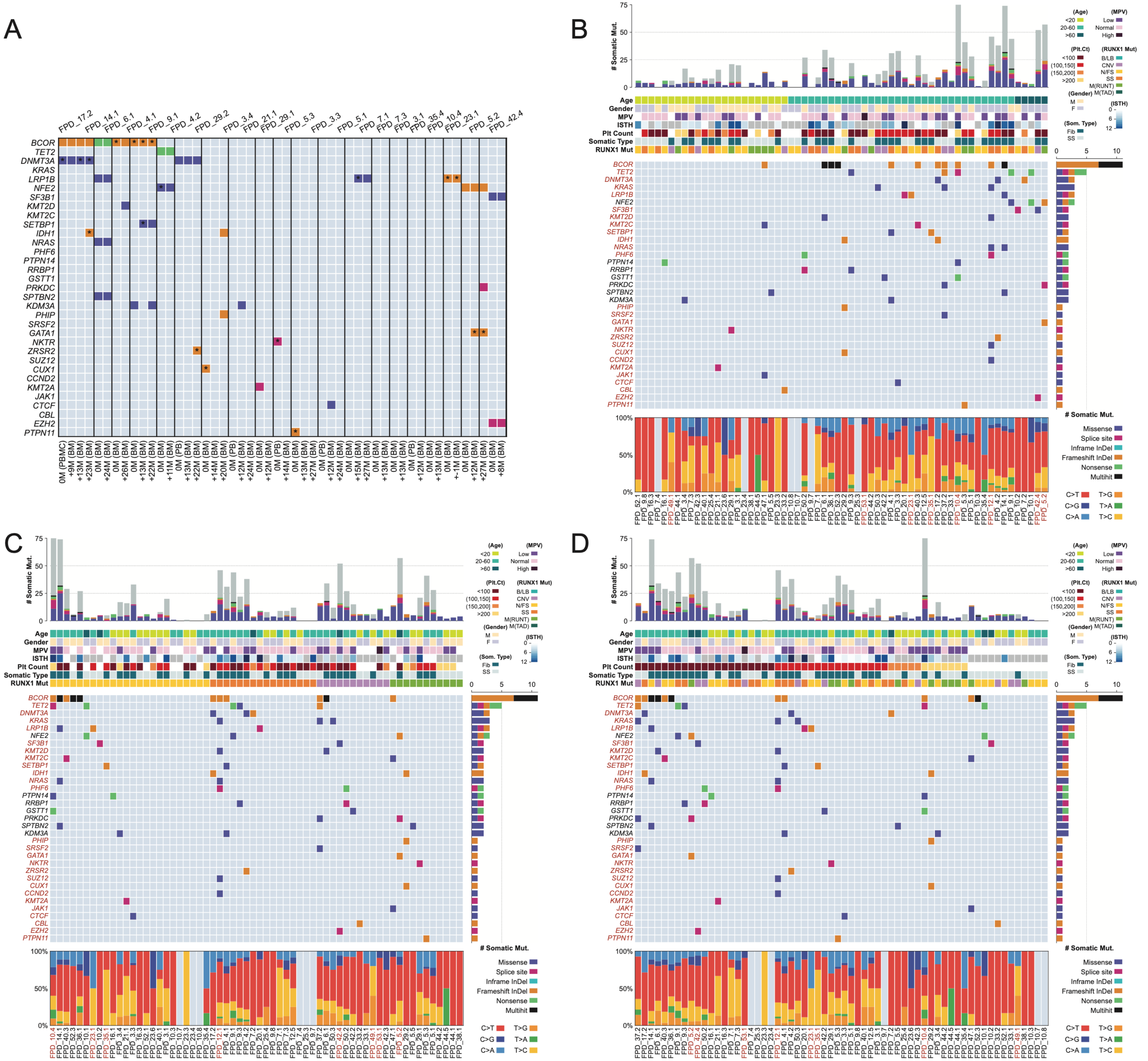

A
B
C
D
BCOR
TET2
DNMT3A
KRAS
LRP1B
NFE2
SF3B1
KMT2D
KMT2C
SETBP1
IDH1
NRAS
PHF6
PTPN14
RRBP1
GSTT1
PRKDC
SPTBN2
KDM3A
PHIP
SRSF2
GATA1
NKTR
ZRSR2
SUZ12
CUX1
CCND2
KMT2A
JAK1
CTCF
CBL
EZH2
PTPN11
### Somatic Mut.
Age
Gender
MPV
ISTH
Plt Count
Somatic Type
RUNX1 Mut
(Age)
<20
20-60
>60
(MPV)
Low
Normal
High
(Plt.Ct)
<100
(100,150]
(150,200]
>200
(RUNX1 Mut)
B/LB
CNV
N/FS
SS
M(RUNT)
M(TAD)
(Gender)
M
F
(ISTH)
(Som. Type)
Fib
SS
Missense
Splice site
Inframe InDel
Frameshift InDel
Nonsense
Multihit
C>T
T>G
C>G
T>A
C>A
T>C

Figure S4 (cont.)

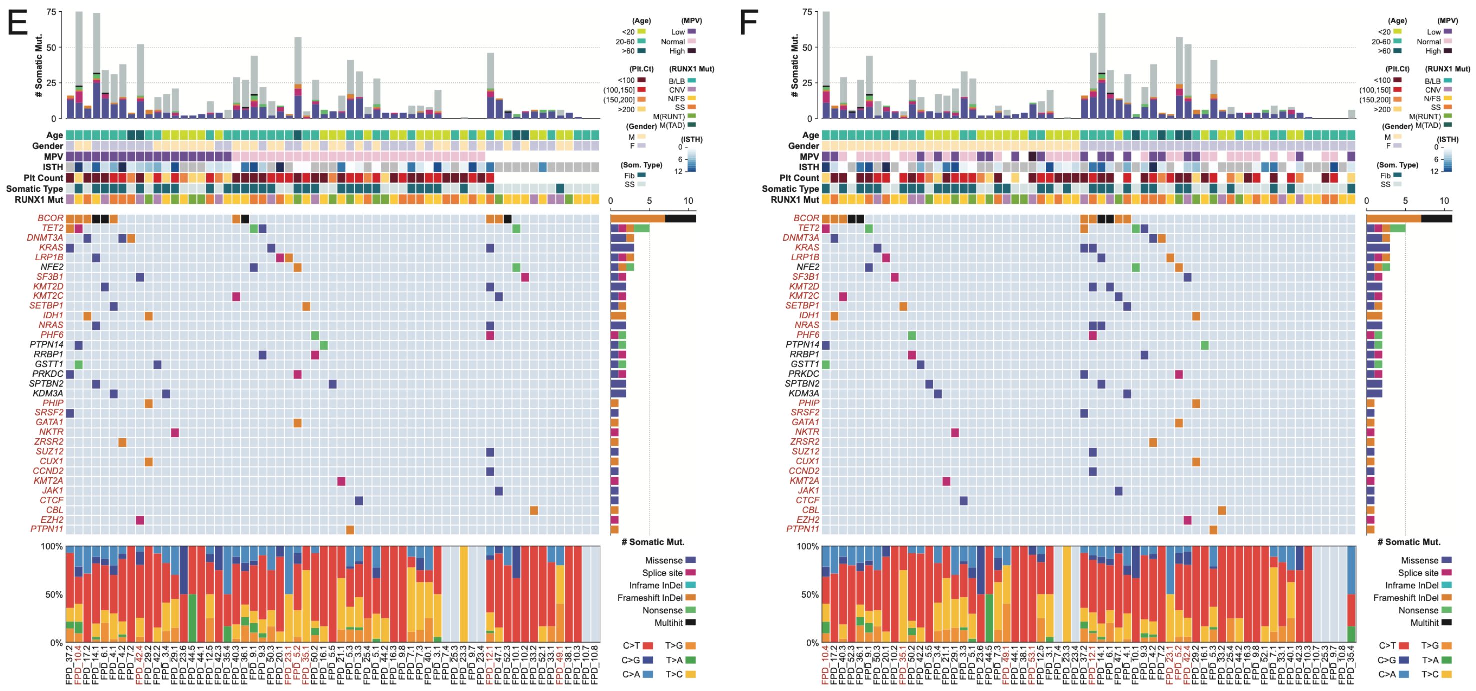


Supplemental Figure 4. **Somatic mutation comut plots related to Figure 3A** (A) Breakdown comut plot of patients with samples from multiple timepoints. Star signs indicate the mutation VAF<3%. Comut plot as shown in Fig 3A, being sorted by age (B), *RUNX1* mutation types (C), platelet count (D), mean platelet volume (MPV) (E) , and gender (F).

Figure S5

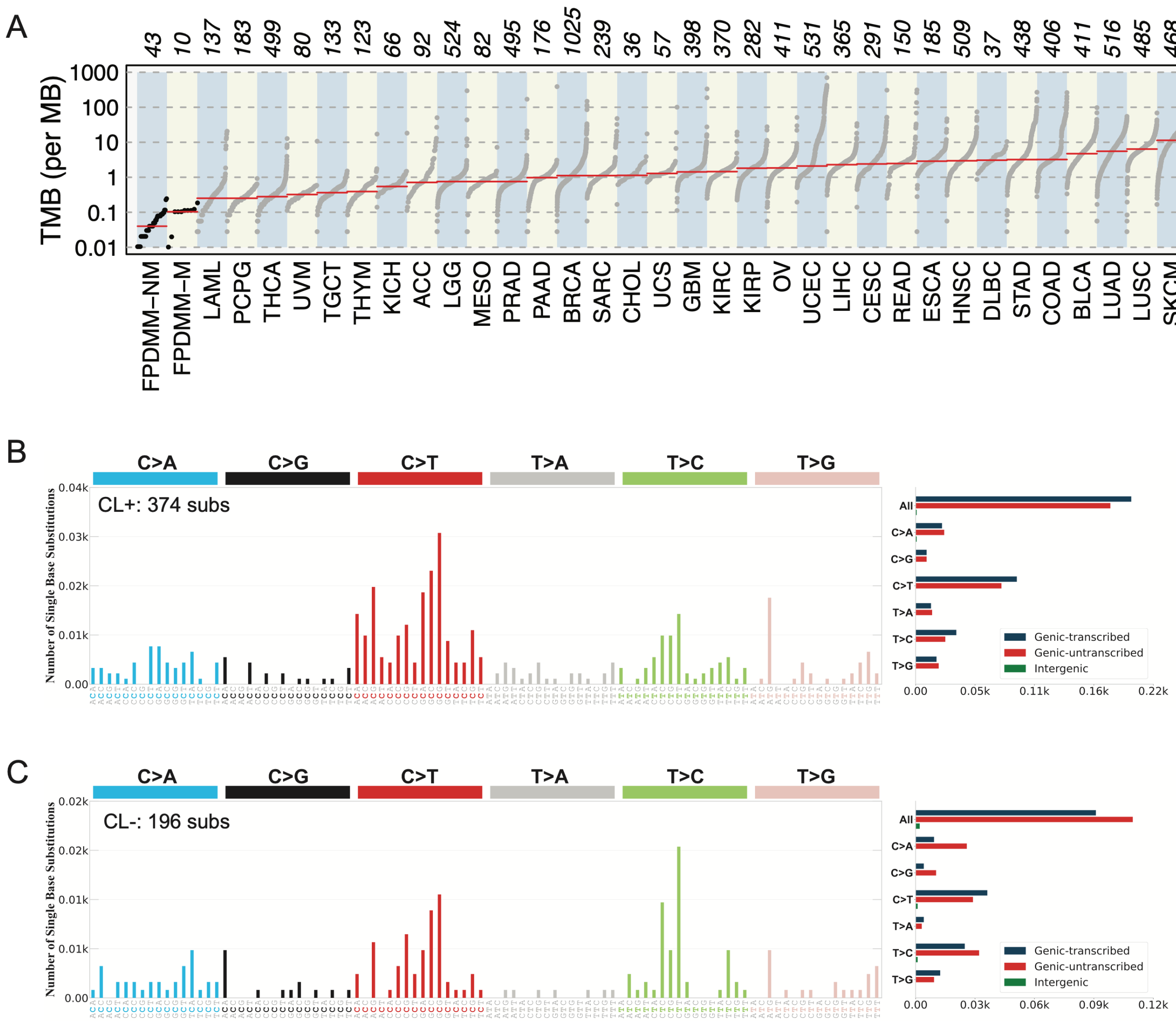
A
B
C
TMB (per MB)
FPDMM–NM
FPDMM–M
LAML
PCPG
THCA
UVM
TGCT
THYM
KICH
ACC
LGG
MESO
PRAD
PAAD
BRCA
SARC
CHOL
UCS
GBM
KIRC
KIRP
OV
UCEC
LIHC
CESC
READ
ESCA
HNSC
DLBC
STAD
COAD
BLCA
LUAD
LUSC
SKCM
CL+: 374 subs
CL-: 196 subs
Number of Single Base Substitutions
C>A
C>G
C>T
T>A
T>C
T>G
All
Genic-transcribed
Genic-untranscribed
Intergenic

**Supplemental Figure 5. Characterizations of somatic mutations in the NIH FPDMM cohort**. (A) Somatic mutation burdens of non-malignant, and malignant FPDMM patients, compared to the somatic mutation burdens of other cancer types from TCGA project. TMB: tumor mutational burden. Somatic mutation signatures of non-malignant FPDMM patients who carry somatic mutations in CHIP genes and AML driver genes (B) and of those who do not (C).

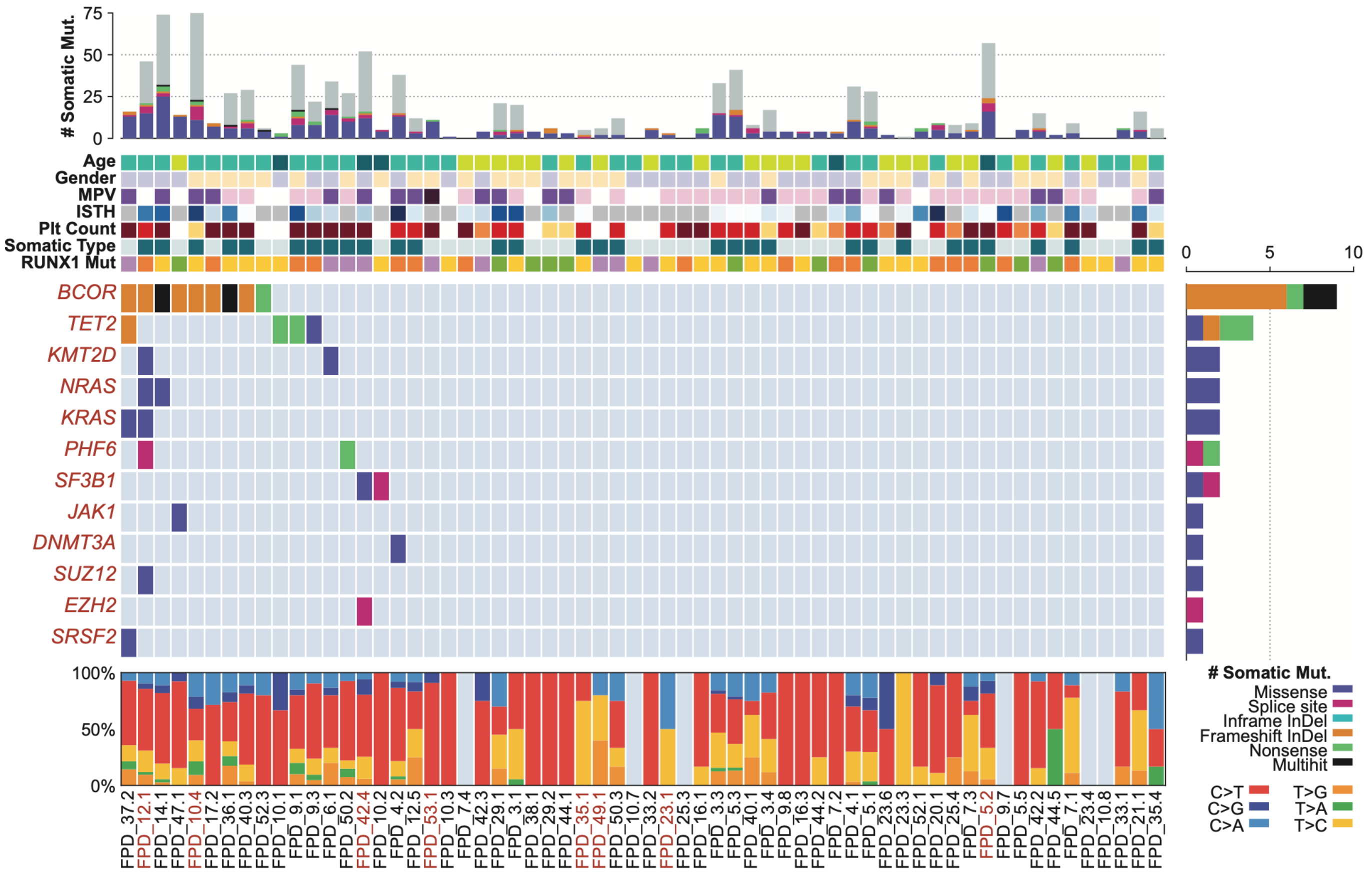


Supplemental Figure 6. Somatic mutation comut plots of variants detected in 74 CHIP gene list and with allele frequency (VAF) >5%.

#### Figure S7

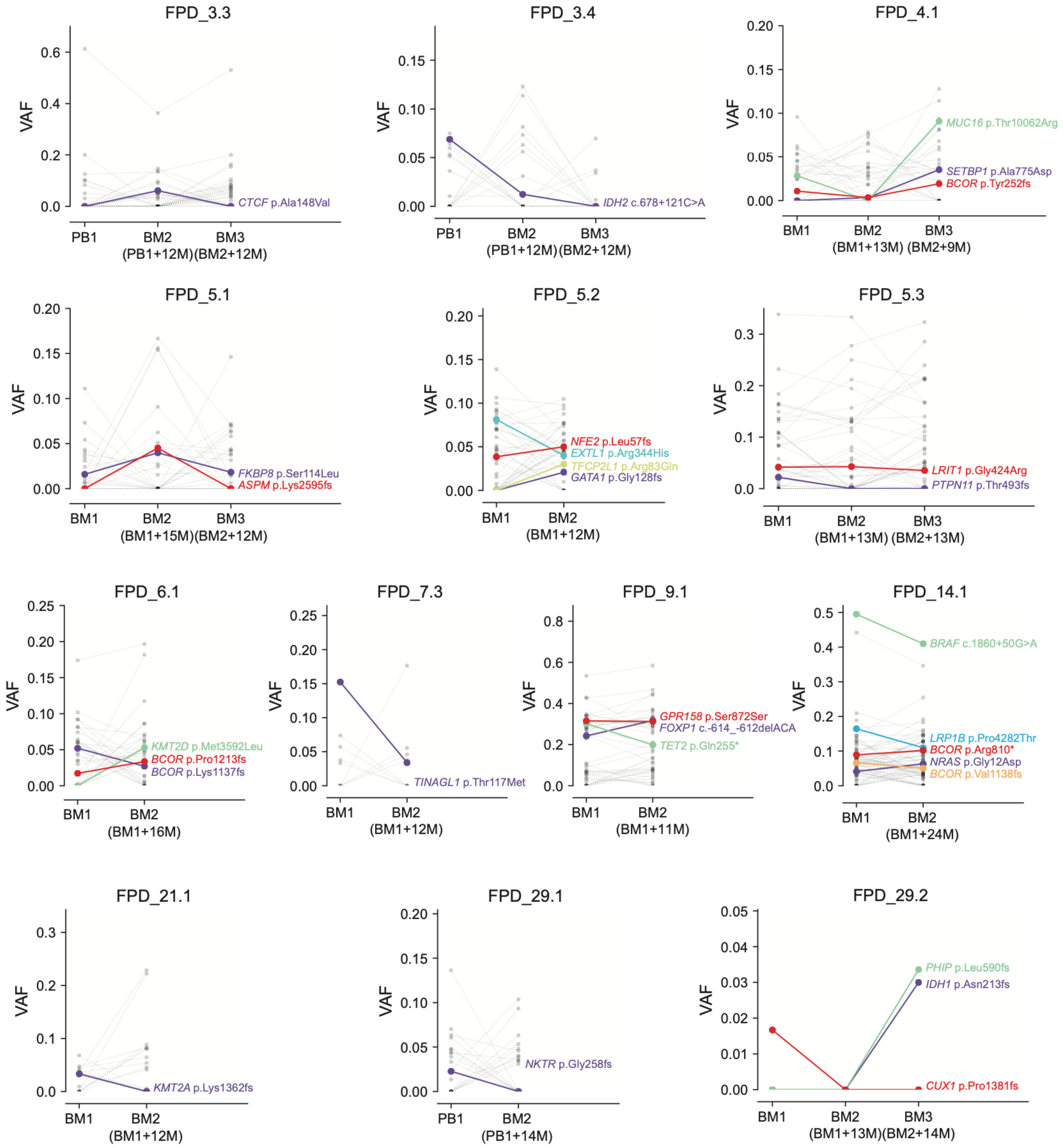

FPD_3.3
VAF
CTCF p.Ala148Val
PB1
BM2 (PB1+12M)
BM3 (BM2+12M)
FPD_3.4
IDH2 c.678+121C>A
PB1
BM2 (PB1+12M)
BM3 (BM2+12M)
FPD_4.1
MUC16 p.Thr10062Arg
SETBP1 p.Ala775Asp
BCOR p.Tyr252fs
BM1
BM2 (BM1+13M)
BM3 (BM2+9M)
FPD_5.1
FKBP8 p.Ser114Leu
ASPM p.Lys2595fs
BM1
BM2 (BM1+15M)
BM3 (BM2+12M)
FPD_5.2
NFE2 p.Leu57fs
EXTL1 p.Arg344His
TFCP2L1 p.Arg83Gln
GATA1 p.Gly128fs
BM1
BM2 (BM1+12M)
FPD_5.3
LRIT1 p.Gly424Arg
PTPN11 p.Thr493fs
BM1
BM2 (BM1+13M)
BM3 (BM2+13M)
FPD_6.1
KMT2D p.Met3592Leu
BCOR p.Pro1213fs
BCOR p.Lys1137fs
BM1
BM2 (BM1+16M)
FPD_7.3
TINAGL1 p.Thr117Met
BM1
BM2 (BM1+12M)
FPD_9.1
GPR158 p.Ser872Ser
FOXP1 c.-614_-612delACA
TET2 p.Gln255*
BM1
BM2 (BM1+11M)
FPD_14.1
BRAF c.1860+50G>A
LRP1B p.Pro4282Thr
BCOR p.Arg810*
NRAS p.Gly12Asp
BCOR p.Val1138fs
BM1
BM2 (BM1+24M)
FPD_21.1
KMT2A p.Lys1362fs
BM1
BM2 (BM1+12M)
FPD_29.1
NKTR p.Gly258fs
PB1
BM2 (PB1+14M)
FPD_29.2
PHIP p.Leu590fs
IDH1 p.Asn213fs
CUX1 p.Pro1381fs
BM1
BM2 (BM1+13M)
BM3 (BM2+14M)

Supplemental Figure 7. Somatic mutation VAF changes across samples collected in different timepoints from 13 non-malignant FPDMM patients. Colored dots and lines show mutations in CL and leukemia genes. Grey dots and lines show somatic mutations in other genes.

Figure S8

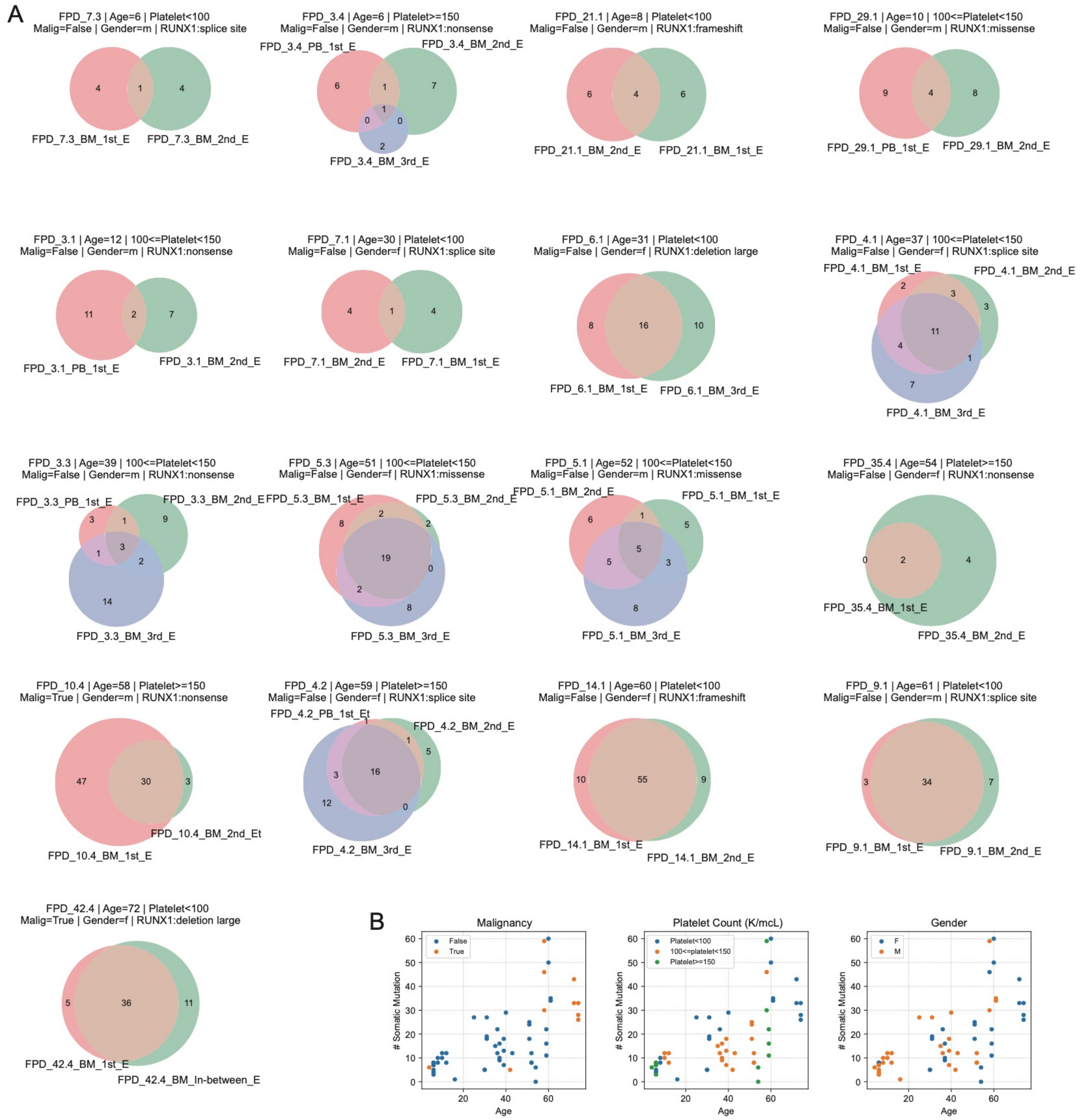
A
FPD_7.3 | Age=6 | Platelet<100
Malig=False | Gender=m | RUNX1:splice site
4
1
4
FPD_7.3_BM_1st_E
FPD_7.3_BM_2nd_E
FPD_3.4 | Age=6 | Platelet>=150
Malig=False | Gender=m | RUNX1:nonsense
FPD_3.4_PB_1st_E
FPD_3.4_BM_2nd_E
6
1
7
1
0
0
2
FPD_3.4_BM_3rd_E
FPD_21.1 | Age=8 | Platelet<100
Malig=False | Gender=m | RUNX1:frameshift
6
4
6
FPD_21.1_BM_2nd_E
FPD_21.1_BM_1st_E
FPD_29.1 | Age=10 | 100<=Platelet<150
Malig=False | Gender=m | RUNX1:missense
9
4
8
FPD_29.1_PB_1st_E
FPD_29.1_BM_2nd_E
FPD_3.1 | Age=12 | 100<=Platelet<150
Malig=False | Gender=m | RUNX1:nonsense
11
2
7
FPD_3.1_PB_1st_E
FPD_3.1_BM_2nd_E
FPD_7.1 | Age=30 | Platelet<100
Malig=False | Gender=f | RUNX1:splice site
4
1
4
FPD_7.1_BM_2nd_E
FPD_7.1_BM_1st_E
FPD_6.1 | Age=31 | Platelet<100
Malig=False | Gender=f | RUNX1:deletion large
8
16
10
FPD_6.1_BM_1st_E
FPD_6.1_BM_3rd_E
FPD_4.1 | Age=37 | 100<=Platelet<150
Malig=False | Gender=f | RUNX1:splice site
FPD_4.1_BM_1st_E
FPD_4.1_BM_2nd_E
2
3
3
11
4
1
7
FPD_4.1_BM_3rd_E
FPD_3.3 | Age=39 | 100<=Platelet<150
Malig=False | Gender=m | RUNX1:nonsense
FPD_3.3_PB_1st_E
FPD_3.3_BM_2nd_E
3
1
9
1
3
2
14
FPD_3.3_BM_3rd_E
FPD_5.3 | Age=51 | 100<=Platelet<150
Malig=False | Gender=f | RUNX1:missense
FPD_5.3_BM_1st_E
FPD_5.3_BM_2nd_E
8
2
2
19
0
2
8
FPD_5.3_BM_3rd_E
FPD_5.1 | Age=52 | 100<=Platelet<150
Malig=False | Gender=m | RUNX1:missense
FPD_5.1_BM_2nd_E
FPD_5.1_BM_1st_E
6
1
5
5
5
3
8
FPD_5.1_BM_3rd_E
FPD_35.4 | Age=54 | Platelet>=150
Malig=False | Gender=f | RUNX1:nonsense
0
2
4
FPD_35.4_BM_1st_E
FPD_35.4_BM_2nd_E
FPD_10.4 | Age=58 | Platelet>=150
Malig=True | Gender=m | RUNX1:nonsense
47
30
3
FPD_10.4_BM_2nd_Et
FPD_10.4_BM_1st_E
FPD_4.2 | Age=59 | Platelet>=150
Malig=False | Gender=f | RUNX1:splice site
FPD_4.2_PB_1st_Et
FPD_4.2_BM_2nd_E
1
1
5
3
16
12
0
FPD_4.2_BM_3rd_E
FPD_14.1 | Age=60 | Platelet<100
Malig=False | Gender=f | RUNX1:frameshift
10
55
9
FPD_14.1_BM_1st_E
FPD_14.1_BM_2nd_E
FPD_9.1 | Age=61 | Platelet<100
Malig=False | Gender=m | RUNX1:splice site
3
34
7
FPD_9.1_BM_1st_E
FPD_9.1_BM_2nd_E
FPD_42.4 | Age=72 | Platelet<100
Malig=True | Gender=f | RUNX1:deletion large
5
36
11
FPD_42.4_BM_1st_E
FPD_42.4_BM_In-between_E
B
Malignancy
False
True
### Somatic Mutation
Age
Platelet Count (K/mcL)
Platelet<100
100<=platelet<150
Platelet>=150
### Somatic Mutation
Age
Gender
F
M
### Somatic Mutation
Age
0
10
20
30
40
50
60
20
40
60

Supplemental Figure 8. Somatic mutations detected at more than one timepoints. (A) Venn plots of somatic mutations identified at different timepoints. Numbers indicate mutations detected at one or more timepoints. (B) Numbers of somatic mutations correlating with patient age, malignancy status, platelet count, and gender. Each dot represent one unique patient (?).

Figure S9

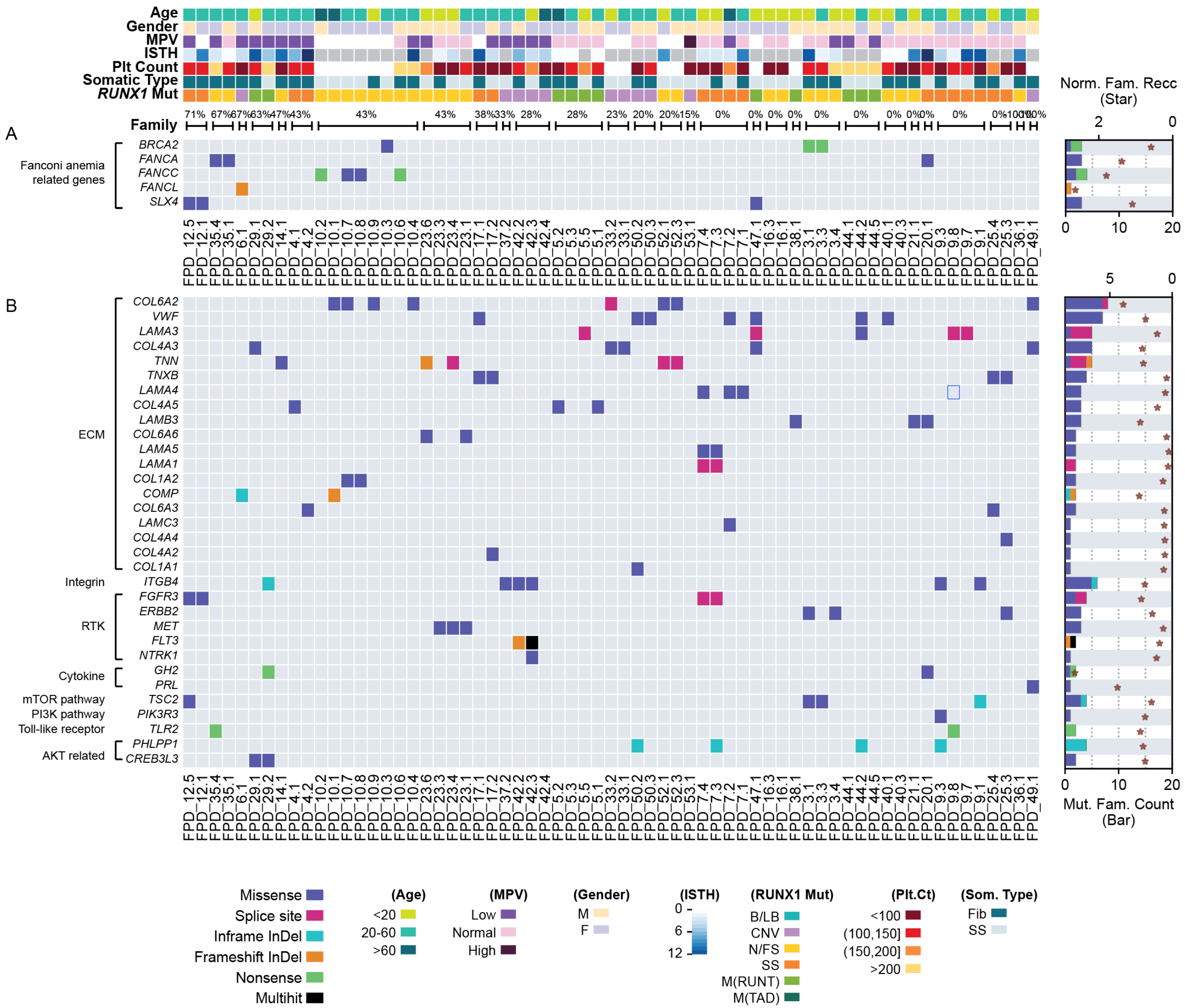


Supplemental Figure 9. **Germline variants in genes related to (A) Fanconi anemia, and (B) genes in the RTK-RAS-PI3K pathway.** The design and color codes for this figure are similar to those in Fig 3A, S5, and S6.
